# Strain-resolved de-novo metagenomic assembly of viral genomes and microbial 16S rRNAs

**DOI:** 10.1101/2024.03.29.587318

**Authors:** Annika Jochheim, Florian A. Jochheim, Alexandra Kolodyazhnaya, Étienne Morice, Martin Steinegger, Johannes Söding

## Abstract

**Background:** Metagenomics is a powerful approach to study environmental and human-associated microbial communities and, in particular, the role of viruses in shaping them. Viral genomes are challenging to assemble from metagenomic samples due to their genomic diversity caused by high mutation rates. In the standard de Bruijn graph assemblers, this genomic diversity leads to complex *k*-mer assembly graphs with a plethora of loops and bulges that are challenging to resolve into strains or haplotypes because variants more than the *k*-mer size apart cannot be phased. In contrast, overlap assemblers can phase variants as long as they are covered by a single read.

**Results:** Here, we present PenguiN, a software for strain resolved assembly of viral DNA and RNA genomes and bacterial 16S rRNA from shotgun metagenomics. Its exhaustive detection of all read overlaps in linear time combined with a Bayesian model to select strain-resolved extensions allow it to assemble longer viral contigs, less fragmented genomes, and more strains than existing assembly tools, on both real and simulated datasets. We show a 3-40-fold increase in complete viral genomes and a 6-fold increase in bacterial 16S rRNA genes.

**Conclusion:** PenguiN is the first overlap-based assembler for viral genome and 16S rRNA assembly from large and complex metagenomic datasets, which we hope will facilitate studying the key roles of viruses in microbial communities.

## 1 Background

In shotgun metagenomics, DNA from environmental samples is directly sequenced, usually with short-read sequencing techniques. Millions to billions of short reads per sample are assembled into contigs, which are then binned into hundreds of different metagenome-assembled genomes. This approach circumvents the need to cultivate microbes in the lab, which requires to painstakingly search for suitable cultivation conditions for each microbial species. The ease of accessing the composition and gene-encoded functions of microbes with these cultivation-independent techniques have therefore greatly propelled progress in environmental microbiology [1] and particularly in medical research, where ever tighter links are being discovered between our gut-associated microbiomes and the development and homeostasis of our immune system, our metabolic health, and the normal functioning of our brains [2, 3, 4].

Viruses infecting bacteria or archaea (called phages) have recently moved into the limelight, as their central roles in shaping environmental and gut microbiomes are becoming clearer. Phages are particularly difficult to study in the lab because their prokaryotic hosts are often difficult to cultivate themselves and, on top, propagating viruses in the lab requires finding conditions that may well be distinct from those used to cultivate their hosts [5]. Therefore, viral metagenomics has greatly accelerated the study of phages and their roles in shaping environmental, animal- and plant-associated microbial ecosystems, such as soil [6, 7, 8], aquatic environments [9, 10, 11, 12, 13, 14], or the human gut [15, 16, 17, 18]. These and other studies indicate that phages have a huge impact on their host communities, by shaping their compositions and dynamics and driving bacterial diversity [19, 20]. However, only a minute sliver of the world’s virome has so far been discovered [21].

Virus genomes can be retrieved from datasets enriched for virus particles (viral metagenomes) or from bulk metagenomes, including both virus particles and microbial cells [22, 5]. Assembling viral genomes from metagenomic data is challenging. Viruses have small sizes and therefore often represent only a minor fraction of the reads, whereas the background contamination from bacterial or eukaryotic hosts can be high [23, 24]. The chief difficulty stems from the high microdiversity and strain heterogeneity resulting from an error prone replication process, the lack of repair mechanisms, and the frequent exchange of genomic segments [25, 26].

Genomic assembly follows one of two approaches [27]: In overlap assembly, one computes the overlap alignments between all reads, links reads with sufficient overlap and similarity in their overlapping regions, and constructs contiguous assembled sequences (contigs) from chains of linked reads. This approach dominated until short read sequencers producing millions of reads became popular and the quadratic runtime for computing all-versus-all alignments became prohibitive. In de Bruijn graph assembly, one constructs a graph in which each node represents a *k*-nucleotide subsequence (*k*-mer) and edges are drawn between successive *k*-mers in a read. Contigs are constructed from paths through this de Bruijn graph. Because no all-versus-all alignment of reads is required, the runtime scales only linearly in the number of input reads. For this reason, modern metagenomics assemblers (except Plass [28]) employ de Bruijn graphs [29, 30, 31].

However, this efficiency gain comes with the downside that genomes of closely related strains cannot be resolved. Beyond the distance of the maximum *k*-mer size, which is typically around 55, the information whether two variants occurred in the same read is lost when reducing the reads to a de Bruijn graph (Fig. **1**B). Genomes with average nucleotide identity (ANI) ≥ 95% have many identical stretches longer than the typical *k*-mer size. The *k*-mer size is limited by the quickly exploding complexity of the de Bruijn graph for increasing *k* due to intra-strain microdiversity in microbial populations and sequencing errors. Therefore, most de Bruijn graph assemblers only attempt to assemble consensus genomes instead of strain-resolved genomes.

**Figure 1:**
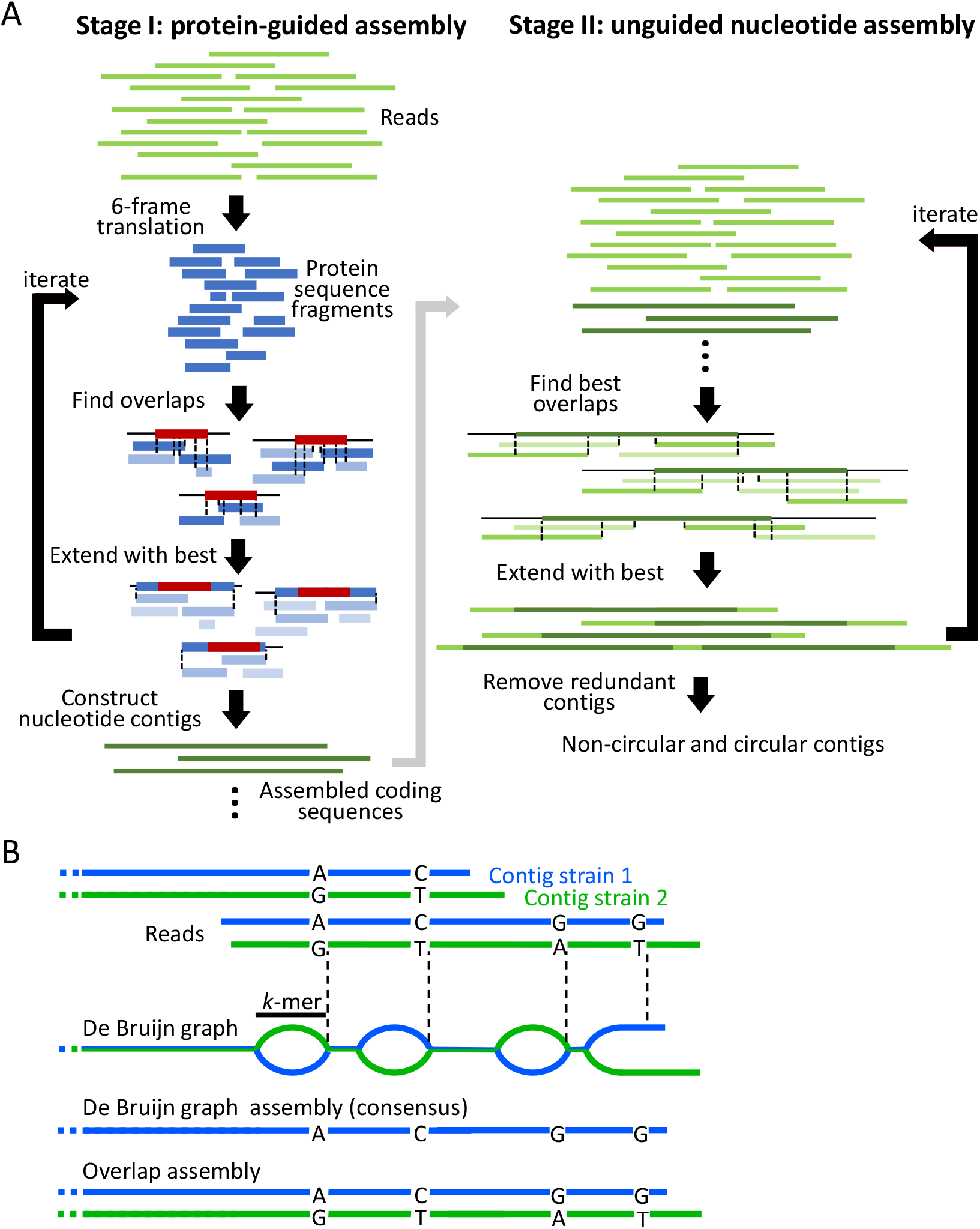
PenguiN overview. (**A**) PenguiN proceeds in two stages. During stage I, PenguiN assembles six-frame translated reads (blue). All sequences sharing a *k*-mer are grouped together. In each *k*-mer group, all sequences are aligned to the longest one (red), and the best-matching sequences are used to extend the red sequence. Penguin keeps track of the underlying nucleotide sequences, resulting in assembled coding sequences (dark green). During stage II, the coding sequences are combined with nucleotide reads (light green). In both stages, sequences are iteratively extended by computing overlap alignments (black dotted lines) in linear time, ranking possible extensions by their quality, and extending upstream and downstream with the best sequence. (**B**) De Bruijn graph assemblers cannot resolve strains across conserved regions longer than their *k*-mer length (black bar). In contrast, overlap assemblers like PenguiN can link strain assemblies correctly across all conserved regions shorter than the (paired) read length.

Viral populations with their high microdiversity and strain heterogeneity pose formidable challenges to assembly of viral genomes from metagenomic data. The resulting complex graph structures lead to fragmented assemblies and make the reconstruction of strain-resolved genomes even harder [32]. State-of-the-art metagenomic viral assemblers as metaviralSPAdes [33], rnaviralSPAdes [34], and Phables [35] employ various heuristics to simplify the graph structure, for example by removing tips, bubbles, and bulges with low coverage. To resolve strains, Phables, rnaviralSPAdes, and Haploflow [36] employ flow decomposition: They find a set of paths and read coverages that would add up to approximately the read coverages observed in the de Bruijn graph. Unfortunately, flow decomposition has less utility for virus genomes due to their uneven read coverage, particularly for RNA phages, for which the reverse transcription step introduces strong sequence biases [37].

Several tools have been developed for the assembly of viral genomes in patient samples, which typically contain a small number of highly similar strains. Haplotype assemblers like Haploflow, SAVAGE [38], and ViPRA-Haplo [39] aim to reconstruct strain-resolved genomes, while IVA [40] and VICUNA [41] reconstruct consensus genomes. All except Haploflow are overlap assemblers, which make use of the co-occurrence of mutations within reads. However, they were not designed to analyse complex metagenomic dataset and also tend to be much slower than de Bruijn graph assemblers.

Assembly of 16S rRNA genes from metagenomic data is of particular interest because of their use as marker genes in the identification, quantification, and phylogenetic analysis of prokaryotes. Assembling 16S rRNA genes is a problem analogous to strain-resolved viral genome assembly. Like genomes of related viral strains, 16S rRNA genes have highly conserved regions interspersed with regions of higher variation. The highly conserved regions are longer than the typical *k*-mer length, and hence de Bruijn graph assemblers cannot know how to connect these variable regions across the conserved ones. This results in highly branched de Bruijn graphs, making the reconstruction of individual 16S rRNA gene sequences hard [42, 43]. Usually, either 16S sequences of several species are merged into consensus sequences or assemblies remain very fragmented [44].

Here, we present PenguiN, a de-novo metagenomic overlap assembler for viral genomes and 16S rRNA genes. PenguiN iteratively extends reads and growing contigs by computing all overlap alignments between them and selecting the extensions that are most likely to be from the same strain, based on a novel Bayesian statistical model. Similar to Plass [28], our protein-level assembler, PenguiN achieves fast speeds and a linear run time complexity by applying Linclust [45], the first algorithm to cluster protein and nucleotide sequences in linear time. To gain robustness with respect to the high level of microdiversity in viral genomes, PenguiN guides its nucleotide assembly by evaluating the protein sequences when merging coding sequences. On simulated and real complex metagenomic short read datasets, PenguiN assembled several times more viral genomes and 16S RNAs at similar error rates than nine state-of-the-art metagenomic or viral assemblers.

## 2 Results

### Overview of PenguiN

PenguiN is an overlap assembler that uses two largely analogous stages. In stage I (Fig. **1**A left), it assembles six-frame translated reads into proteins, while also co-assembling the corresponding nucleotide coding sequences. In stage II (Fig. **1**A right), it links the contigs covering coding regions obtained from stage I across non-coding regions. PenguiN follows a greedy iterative assembly scheme. Like our protein-level assembler Plass [28], it uses the Linclust algorithm [45] to find statistically significant overlaps in linear time and extends the sequences iteratively. To avoid chimeric extensions, for each center sequence, we select from several possible extensions the one with the highest Bayesian posterior probability to belong to the same genome (see Methods). When overlap lengths differ, this scheme improves upon the Plass extension strategy of choosing the overlap with the highest sequence identity. Both stages perform multiple iterations of overlap computation and extension. In addition, in stage II we detect circular sequences after each extension (see Methods), because plasmids and many viral genomes are circular. In contrast to the graph-based assemblers that can detect cycles directly using their graph structure, such detection is not possible in our graph-free assembler. To prevent over-extension for circular genomes or those with long terminal repeats, PenguiN terminates the extension when it detects a cyclic structure, excludes them from further iterations and marks them for the final results. Finally, the resulting contigs are clustered and one representative per cluster is selected to reduce redundancy.

Using full-read overlaps to find the best extensions enables PenguiN to perform strain-level assemblies, a task that standard *k*-mer-based de Bruijn graph assemblers struggle with (Fig. **1**B). Many perfectly conserved regions between two strains will be longer than their *k*-mer length, since strains typically have sequence identities between 90% and 99%. De Bruijn graph assemblers have no way of telling which contig upstream of such a conserved region should be joined to which contig downstream. Therefore, these assemblers either fragment the assembly at the conserved regions to avoid generating chimeric contigs, or they resort to assembling the consensus genome. In contrast, PenguiN can use the full overlap alignment and information about co-occurring mutations to select the best extension and assemble the genomes of strains (Fig. **1**B).

We tested PenguiN on synthetic and real datasets and compared it to state-of-the-art assembly tools, both de Bruijn graph and overlap-based.

### Assembly of an in-silico mixture of HRV genomes

We first generated a very simple synthetic dataset comprised of only three human rhinovirus strains (see Methods). We simulated 2 × 150 bp error-free reads from the three genomes using randomreads.sh from the BBMAP software suite in proportions 4:2:1 with coverages of 200, 100 and 50, respectively. The genomes have average nucleotide identities (ANI) ranging from 92% to 95%, with mismatches mainly due to single-nucleotide polymorphisms (SNPs). We assembled them using PenguiN and nine other assemblers: Megahit [29], metaSPAdes [30], metaviralSPAdes [33], rnaSPAdes [31], rnaviralSPAdes [34], SAVAGE [38], IVA [40], VICUNA [41] and Haploflow [36]. MetaviralSPAdes did not produce any assemblies and was excluded from the subsequent analysis. Assembly quality was assessed using MetaQUAST [46]. The results are shown in Fig. **2** and the MetaQUAST report is available in Additional File 1: Fig. S1.

**Figure 2:**
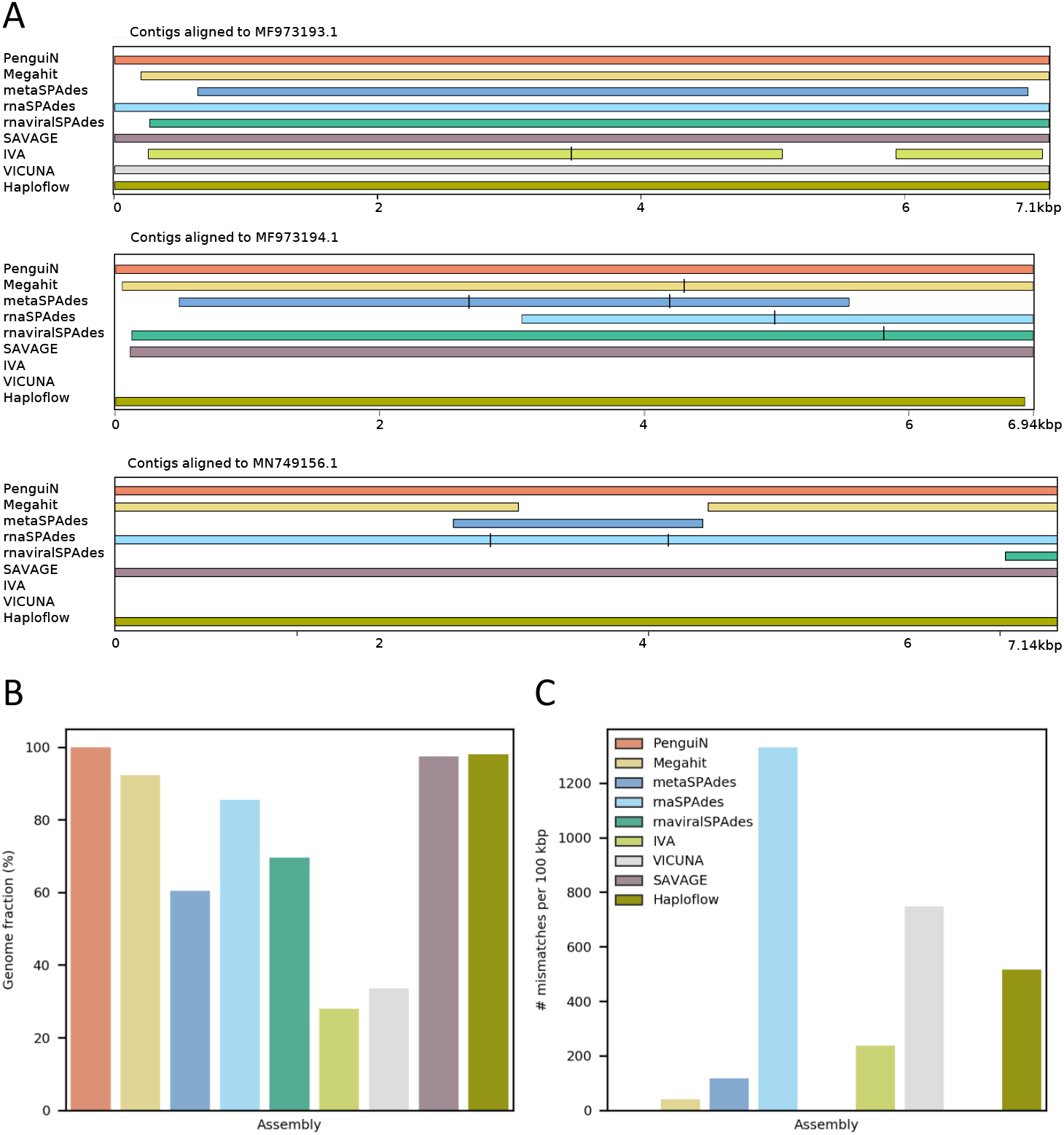
Assembly quality on an in-silico mixture of three HRV genomes. (**A**) Comparison of three genomes assembled by the nine assemblers aligned to the reference. Black bars indicate fragmentation where contigs overlap. (**B**) Fraction of the three genomes covered and (**C**) number of mismatches, computed by MetaQUAST [46].

PenguiN performed well in all metrics. It assembled three contigs, covering almost 100% of all three strain genomes with a single contig each (Fig. **2**A, B). Two further assemblers assembled all three strain genomes to a high degree of completeness: Haploflow (97.98%) and SAVAGE (97.39%). All other assemblers only assembled the most frequent strain completely, while either fragmenting or missing parts of the other two. For example, Megahit recovered only 80% of the strain with the lowest abundance (MN749156.1), metaSPAdes recovered merely 20%, and rnaviralSPAdes only 15%. IVA and VICUNA, which were designed to assemble viral quasispecies, both assembled a consensus genome, that could only be aligned to the most abundant genome by MetaQUAST.

MetaQUAST reported no misassemblies for any of the tools, but half of them generated assemblies with a considerable number of mismatches or indel errors (Fig. **2**C, Additional File 1: Fig. S1), probably due to inter-strain chimeric contigs.

### Performance on highly diverse strain mixtures

Next, to test the ability of assemblers to reconstruct genomes of viral strains from extremely diverse samples, we downloaded 2550 HIV-1 genomes from the NCBI database (see Methods). The HIV-1 genome is approximately 9.7 kbp and has long terminal repeats (LTRs) at both ends, due to which the genome can be both in linear and circular form [47, 48]. We used the circularized forms of all genomes as the ground-truth reference (see Methods), since all linear assemblies can be perfectly mapped to them. The pairwise nucleotide identities among the 2550 genomes are mostly in the range of 85% to 95% (Fig. **3**A). We generated synthetic datasets with 2 × 150 bp error-free reads with overlap lengths uniformly distributed between 40 and 60 nucleotides. To analyse the dependence of the assembly quality on genomic coverage, we generated in total three datasets of reads with mean coverage of each genome set to 1×, 10×, and 100×. Due to the high number and similarity of the genomes, this dataset was particularly challenging to assemble.

**Figure 3:**
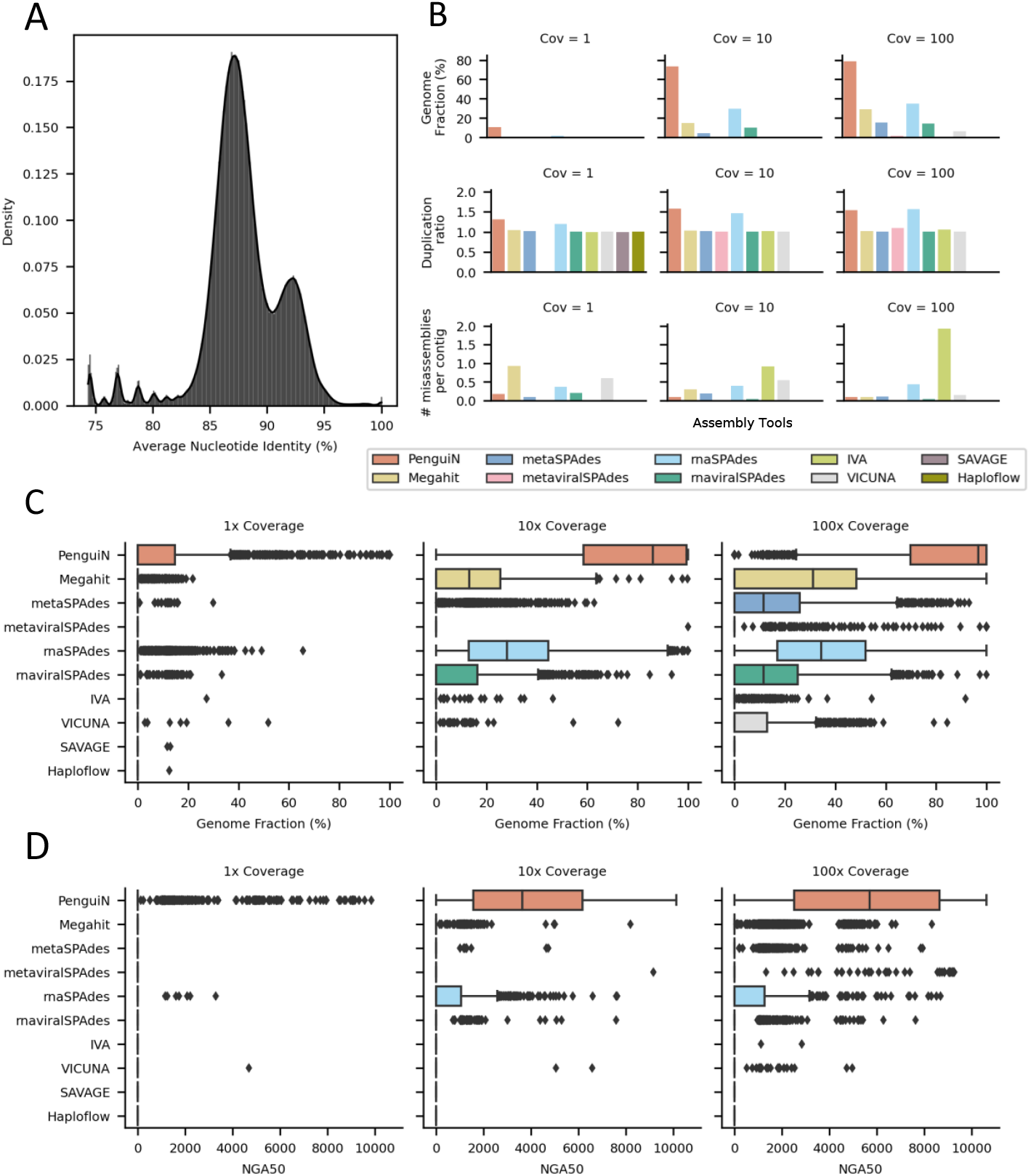
Assembly quality on an in-silico mixture of 2550 HIV-1 genomes on three datasets with an average coverage of each genome of 1×, 10×, and 100×. (**A**) Distribution of pairwise sequence identities among the 2550 HIV-1 genomes. (**B**) Average assembled genome fraction, number of misassemblies per contig, and duplication ratio. (**C**) Distribution of the fractions of each of the 2550 genomes covered by the assemblies. (**D**) Distribution of the NGA50 values. The NGA50 value of a genome assembly is the length for which contigs of that size and longer cover 50% of the genome. The larger the NGA50 value is, the less fragmented is the assembly.

SAVAGE de novo and Haploflow did not finish within 10 days at 10× and 100× coverage and could not be evaluated. MetaviralSPAdes, IVA, and VICUNA produced only very few or no contigs. We again evaluated the assembly quality using MetaQUAST [46]. PenguiN performed best in most metrics (Fig. **3**B-D). The distribution of the fractions of assembled genomes shows that PenguiN assembled several times higher genome fractions across all coverage values than the next best tool, rnaSPAdes (Fig. **3**B). The median assembled genome fractions for PenguiN at 10 and 100 times coverage were 86% and 97%, respectively, compared to 28% and 34% for rnaSPAdes (Fig. **3**C). On the low coverage set, PenguiN was the only tool able to recover genomes with a fraction of more than 90%. The analysis was performed using the MetaQUAST default threshold of 95% sequence identity. In total, PenguiN recovered 27 genomes by more than 90% on the 1× coverage set, 1136 on the 10× coverage set, and 1550 genomes on the 100× coverage set, whereas the next best tools rnaSPAdes and Megahit recovered 0, 24, and 36 and 0, 3 and 11 respectively, corresponding to an approximately 40-fold increase. Further, we also saw that introducing sequencing errors in the simulated reads (see Methods) does not change this picture (see Fig. S2).

PenguiN’s genome assemblies also consist of much longer contigs, as measured by NGA50 (Fig. **3**D). While almost all assemblers can gain from more coverage, most assemblers except for PenguiN either have fragmented assemblies or do not cover the reference genomes with ≥ 50% (counted as NGA50=0). At 10 and 100 times coverage, 50% of the 2550 genomes assembled by PenguiN had NGA50 values above 3.6 kbp and 5.7 kbp, respectively, while the next best tool in this regard, rnaSPAdes, had only 22 and 12 genomes with NGA50 values above these lengths.

Also, PenguiN’s number of misassemblies per contig was the second best on the 1× and 10× coverage dataset, and the third best on 100× coverage dataset. Especially, it was markedly lower than rnaSPAdes, the assembler with the second best completeness. On the flip side, the high completeness of PenguiN assemblies resulted in a higher duplication ratio (between 1.3 and 1.6) than for other tools except rnaSPAdes (Fig. **3**B).

To analyse at what level of sequence similarity various assemblers still distinguish between genomes and assemble them separately without producing chimeras, we assessed how assembled genome fraction (sensitivity) and assembly quality (precision) depend on the stringency of the mapping of contigs to reference genomes. We mapped the contigs assembled by each tool with MMseqs2 [49] to the 2550 reference genomes. We recorded the per-base sensitivity of the assembled contigs, requiring a minimum sequence identity of *x* % for the alignment between the contig and the mapped reference sequence (Fig. **4**A). The sensitivity measures which fraction of the reference genomes are covered by the assemblies. At a very low stringency of 90% sequence identity, all tools achieve close to 100% sensitivity on the 100x coverage dataset. This means that for any reference genome, all tools are able to assemble contigs that together cover each of the 2550 genomes with contigs that have at least 90% sequence identity. The same contig can be mapped to multiple references.

**Figure 4:**
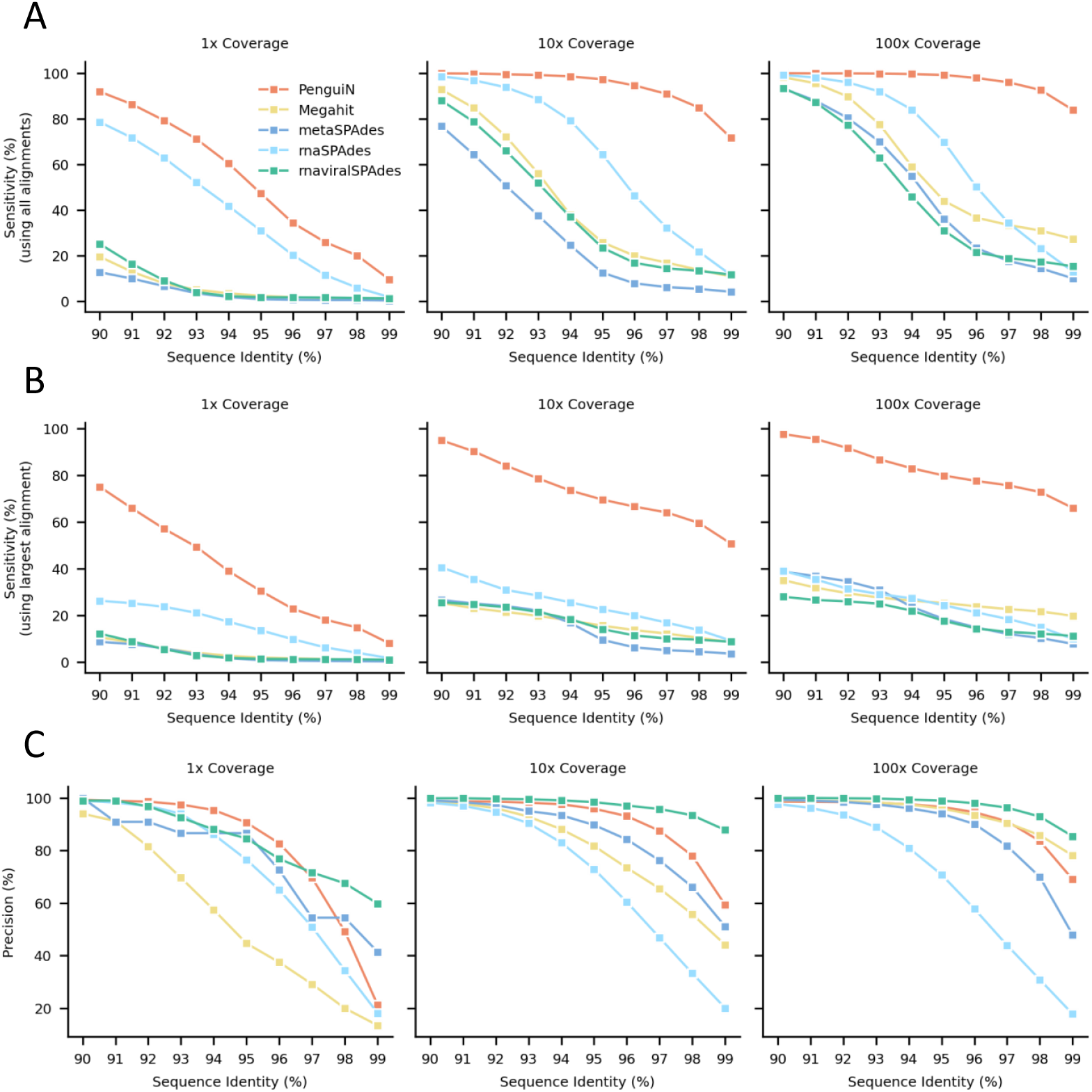
Per-base sensitivity and precision of assemblies generated for the 2550 HIV1 genomes dataset. (**A**) Assembly sensitivity, defined as the fraction of nucleotides in the reference genomes that are aligned to an assembled contig with a sequence identity at least the value on the *x*-axis. (**B**) Fraction of nucleotides of the reference genomes that are aligned to the longest contigs in the assembly that has a sequence identity of at least the value on the *x*-axis. (**C**) Assembly precision is the fraction of assembled nucleotides contained in alignments to a reference genome with sequence identity at least the value on the *x*-axis.

However, PenguiN’s sensitivity decreases much more slowly with increasing stringency than the other tools, demonstrating that it was able to assemble much more of the strain variation in this dataset. At the highest stringency (sequence identity cutoff X=99%), PenguiN correctly assembled approximately 3-6 times more nucleotides than the next best tool rnaSPAdes (1× coverage: 9.6% vs. 1.9%; 10× coverage: 71.6% vs. 11.5%) or Megahit (100× coverage: 83.9% vs. 27.4%). This is in accord with the analysis by MetaQUAST in Fig. **3**B.

When only considering the longest alignment per reference genome instead of all alignments, the difference in sensitivity between PenguiN and the other assemblers becomes even more pronounced across all sequence identity cutoffs (Fig. **4**B). Consistent with the NGA50 analysis, this result reflects the much lower level of fragmentation within PenguiN’s assemblies.

To analyse whether the much longer contigs assembled by PenguiN would lead to a higher fraction of chimeric contigs or misassemblies, we also calculated the per-base precision of the assembled contigs at different stringencies of sequence identity mapping (Fig. **4**C). Precision is the fraction of correctly assembled nucleotides, and one minus precision estimates the error rate, the fraction of assembled nucleotides that cannot be mapped to any reference genome. For all three coverage sets (1×, 10×, 100×) and over the entire range of sequence identities, PenguiN’s precision is close to the second best tool or second best itself, except at coverage 1 and sequence identity ≥ 98%, where

Megahit and rnaviralSPAdes are much more precise but assemble 34 and 50 times fewer nucleotides than PenguiN, respectively. Only rnaviralSPAdes has markedly better precision everywhere, but this comes at the cost of very low sensitivities. At *X* = 99% sequence identity, 21.2%, 59.3%, and 69% of PenguiN’s contigs could still be aligned to a reference genome sequence in the 1×, 10×, and 100× coverage datasets, respectively. These results show that although PenguiN assembles much longer contigs than the other tools, it does not systematically produce more misassemblies.

### Assembly of ssRNA phages from metatranscriptomic samples from activated sludge and aquatic environments

Next, we tested the assemblers on 82 metatranscriptomics samples from activated sludge and aquatic environments, which were recently presented in a study to detect ssRNA phages from these two environments [50]. The challenge in comparing assemblers on real metagenomic data is the lack of a ground truth. Single-stranded RNA (ssRNA) bacteriophages offer a solution, as they occur only in two relatively narrow families, Leviviridae and Levi-like viruses [51]. All of them have a short, positive-stranded genome of 3.5 kbp-4.5 kbp that contains three core proteins, a maturation protein (MP), a coat protein (CP) and an RNA-dependent RNA polymerase (RdRp), as well as a few other facultative, short and much less conserved hypothetical proteins [52, 53, 50]. Since the three core proteins are always present, we could assess the completeness of assembled ssRNA phage genomes by whether they contain full-length versions of the three core proteins.

We assembled the 82 metatranscriptomic samples with PenguiN and the nine other tools and searched the contigs for matches to the profile Hidden Markov Model (HMM) that Callanan et al. [50] constructed by combining HMMs of the three core proteins. Assembly runs were performed on single nodes with two Intel Xeon E5-2640v3 2.6 GHz processors with a total of 16 cores and 128 GB of RAM. As before, the size and complexity of the data set posed problems for multiple assemblers. SAVAGE and IVA did not finish on any sample within 10 days. MetaviralSPAdes assembled only very few contigs with only one hit to the HMM profiles across samples. Haploflow and VICUNA did not finish within 10 days on 5 and 33 samples, respectively, and hence could not be evaluated on these. The runtimes of the assemblers that finished on all samples summed up to 3.7 days for PenguiN, 1.2 days for Megahit, 1.3 days for rnaviralSPAdes, 1.4 days for rnaSPAdes, and 27.8 days for metaSPAdes.

Using the same ssRNA phage detection pipeline and redundancy clustering at 100% as in [50], we identified 29 106 non-redundant ssRNA phage sequences in the PenguiN assemblies across all samples, 4045 of which contain three complete core proteins (“complete genomes”). This number was four times higher than the 1015 complete ssRNA phage genomes previously assembled with rnaSPAdes in [50]. To compare with the other assemblers, we filtered out redundant contig sequences with a stricter sequence identity threshold to avoid counting genome versions differing by only a few mismatches due to sequencing errors or intra-population diversity. We clustered all contigs using MMseqs2 with a maximum sequence identity of 99% and counted only the non-redundant cluster representatives. This reduced the number of sequences containing at least one ssRNA phage protein by a factor of *∼* 2 for all assemblers. In total, this resulted in 1398 “complete genomes” for PenguiN, three times more than for rnaSPAdes and Megahit. Figure **5**A-D compares the number of contigs assembled by PenguiN and the other assemblers at four levels of genome completeness: contigs with a match to a single, two or all three core proteins, and with full-length versions of the three core proteins. At all four completeness levels, PenguiN assembled at least twice as many contigs as the next best tools, Megahit and rnaSPAdes.

**Figure 5:**
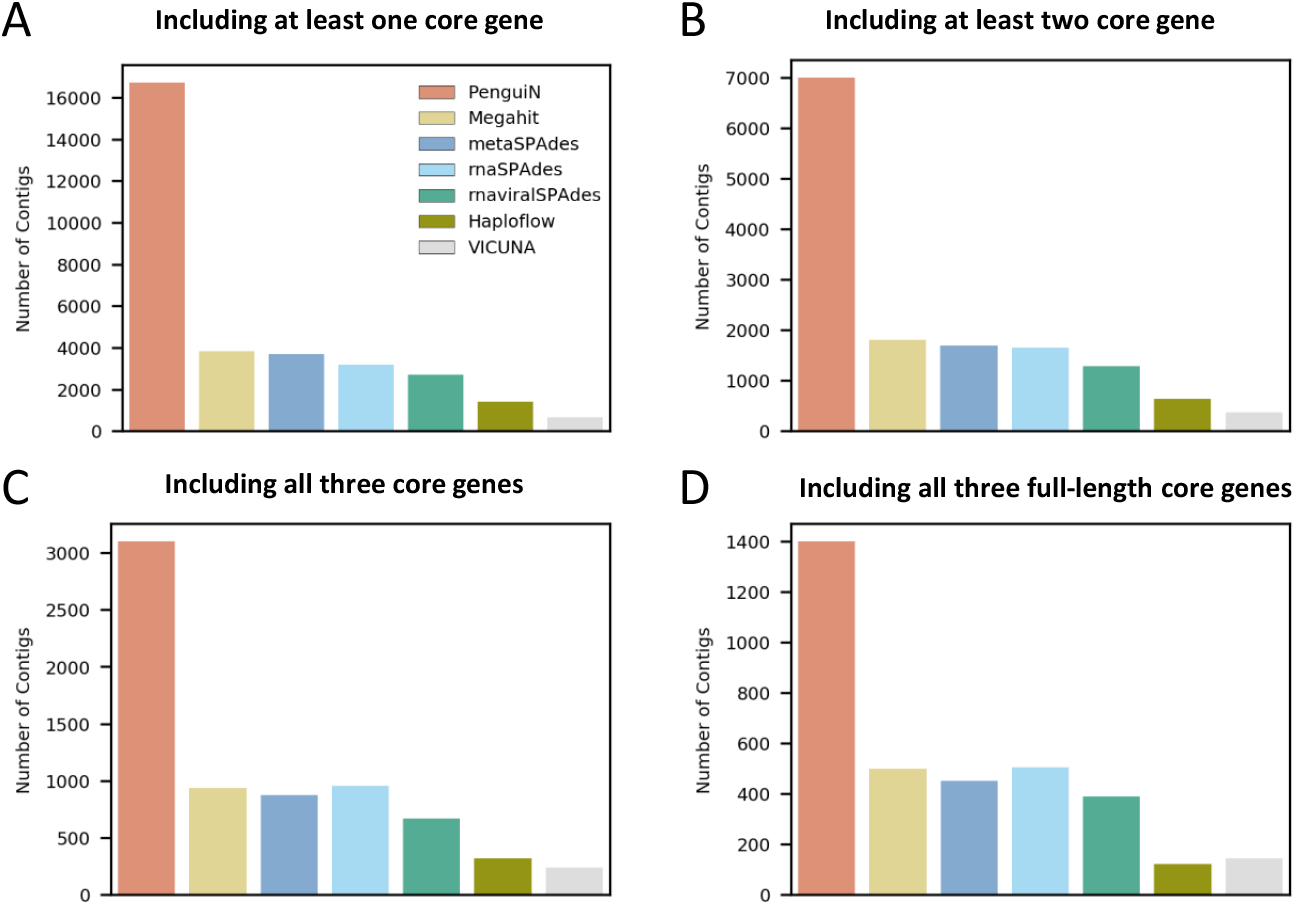
Number of contigs of non-redundant ssRNA phage genomes assembled from 82 environmental metatranscriptomic samples. Profile HMM searches identified contigs containing matches to at least one of the three core proteins of ssRNA phages (MP, CP, RdRp). Contigs were redundancy-filtered at 99% pairwise sequence identity. (**A**) Contigs containing a match to at least one of the three core proteins. (**B**) Contigs containing matches to at least two core proteins. (**C**) Contigs containing matches to all three core proteins. (**D**) Contigs containing full-length matches to all three core proteins (“complete genomes”).

By far the most complete genomes per sample were assembled from activated sludge (samples from Japan, Austria, Illinois), as already reported by [50]. Here, PenguiN assembled 32% more complete genomes than the next best tool, while on the freshwater samples (Mississippi River, and freshwater aquatic samples from Singapore) it yielded a comparable number (Fig. **6**A). On the samples from Lake Mendota, none of the assemblers could reconstruct a complete phage sequence. We then analysed the intersections between each of the assembled sets of complete phage genomes. We aligned the complete contigs of each assembly with the other assemblies and determined for each pair of assemblers (A, B) the contigs from A for which a contig from B exists that has ≥99% identity and covers ≥99% of its residues. We then computed the fraction of assembly A contained in B according to this definition (Methods). The genomes of the three next best assemblers each covered only about one third of the genomes assembled by PenguiN, whereas Penguin covered about two thirds of their genomes (Fig. **6**B). This fraction was quite high considering that PenguiN is an overlap assembler and the other tools are de Bruijn graph assemblers which, among them, also attain fractions around 2/3 despite the similarity of their assembly approaches. The assemblies of Haploflow and VICUNA were even covered by PenguiN with 93.5% and 87.6%, respectively.

**Figure 6:**
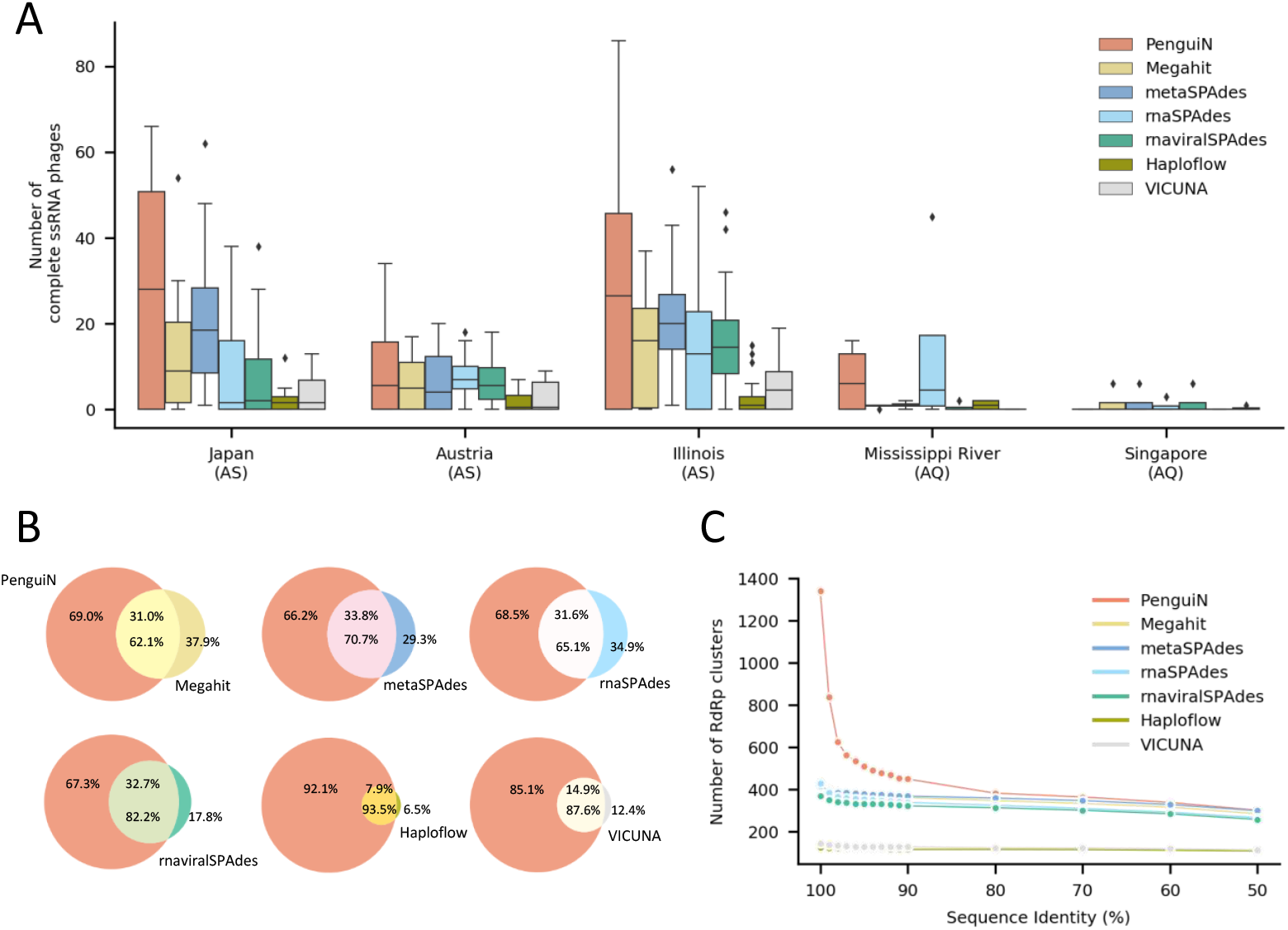
Analysis of assemblies of ssRNA phage genomes from 82 metagenomic samples. (**A**) Distributions of the numbers of complete ssRNA phage genomes for the three sampling locations for activated sludge (AS) and the two environmental freshwater samples that yielded ssRNA genomes (AQ). (**B**) Venn diagrams showing the fractions of contigs of PenguiN covered by contigs of one of the six other tools and contigs of these six tools covered by contigs of PenguiN. For a contig to be covered, at least 99% of its nucleotides must be contained in an alignment with at least 99% sequence identity. (**C**) Number of redundancy-filtered, complete RdRp proteins encoded in the assemblies as a function of stringency of redundancy filtering, the pairwise amino acid sequence identity cut-off.

We asked whether the increased diversity observed in the genomes assembled by PenguiN was due to a higher number of genera, species, or strains. We extracted the RdRp genes from the complete genomes and clustered the RdRp protein sequences at various levels of sequence identity. Using cutoffs for species (80% sequence identity) and genera (50%) found in [50], we saw a similar number of clusters for all assemblies (except VICUNA and Haploflow). However, at higher cutoffs, PenguiN’s assembly showed many more clusters than the other tools, indicating increased strain-level diversity (Fig. **6**C). This was in accordance with our observation on the simulated datasets that Penguin resolved strains where other assemblers merged their genomes.

To check for chimeric assemblies, we considered the genome architecture within PenguiN’s contigs classified as complete ssRNA phage genomes. A simple way to identify chimeric genome assemblies is to count how many of the RNA genomes contain proteins with discordant phylogenies, that is, the co-occurrence of different subgroups of the core genes found in study [50]. Indeed, none of the 1398 complete genomes contained proteins with discordant phylogenies, and all contained exactly one copy of each core gene, except for one contig that contains a second RdRp gene after the first copy (RdRp B) that matches more closely to the other clade (RdRp A) (Additional File 1: Fig. S3). Additionally, the contig lengths of all tools are comparable, with PenguiN producing a somewhat higher fraction of genomes longer than 6 kbp than the other tools (Additional File 1: Fig. S4).

### Assembly of 16S rRNA genes from metatranscriptomes

Amplicon sequencing of 16S rRNA genes is a very popular and cost-effective method to measure the taxonomic composition of prokaryotes in environmental samples [54]. 16S rRNA sequences consist of nine hypervariable regions (V1-V9) connected by eight highly conserved regions. Several of these are nearly perfectly conserved across phyla and longer than typical *k*-mer lengths used in de Bruijn graph assemblers, which makes them challenging to assemble [44]. We therefore used 16S rRNA assembly from metatranscriptomes as another test case for PenguiN. Reference databases for hundreds of thousands of 16S rRNA sequences allow us to verify the correctness of assembled 16S rRNAs. For the assembly of 16S rRNA, we run PenguiN with the same default settings as for viral assembly.

We tested the ability of the assemblers to recover the diverse 16S rRNA genes in metatranscriptomic data. We detected 16S rRNA genes in the assemblies of the 82 metatranscriptomic samples described in the previous section using Barrnap to perform HMM searches using a HMM derived from Rfam (https://github.com/tseemann/barrnap).

PenguiN was able to reconstruct 113 196 non-redundant 16S rRNAs gene fragments that aligned to at least 50% of the Rfam database’s 16S profile HMM model (Fig. **7**A). The second best tool in this category, rnaSPAdes, assembled 18 933 gene fragments at this coverage threshold, *∼* 6 times fewer than PenguiN.

**Figure 7:**
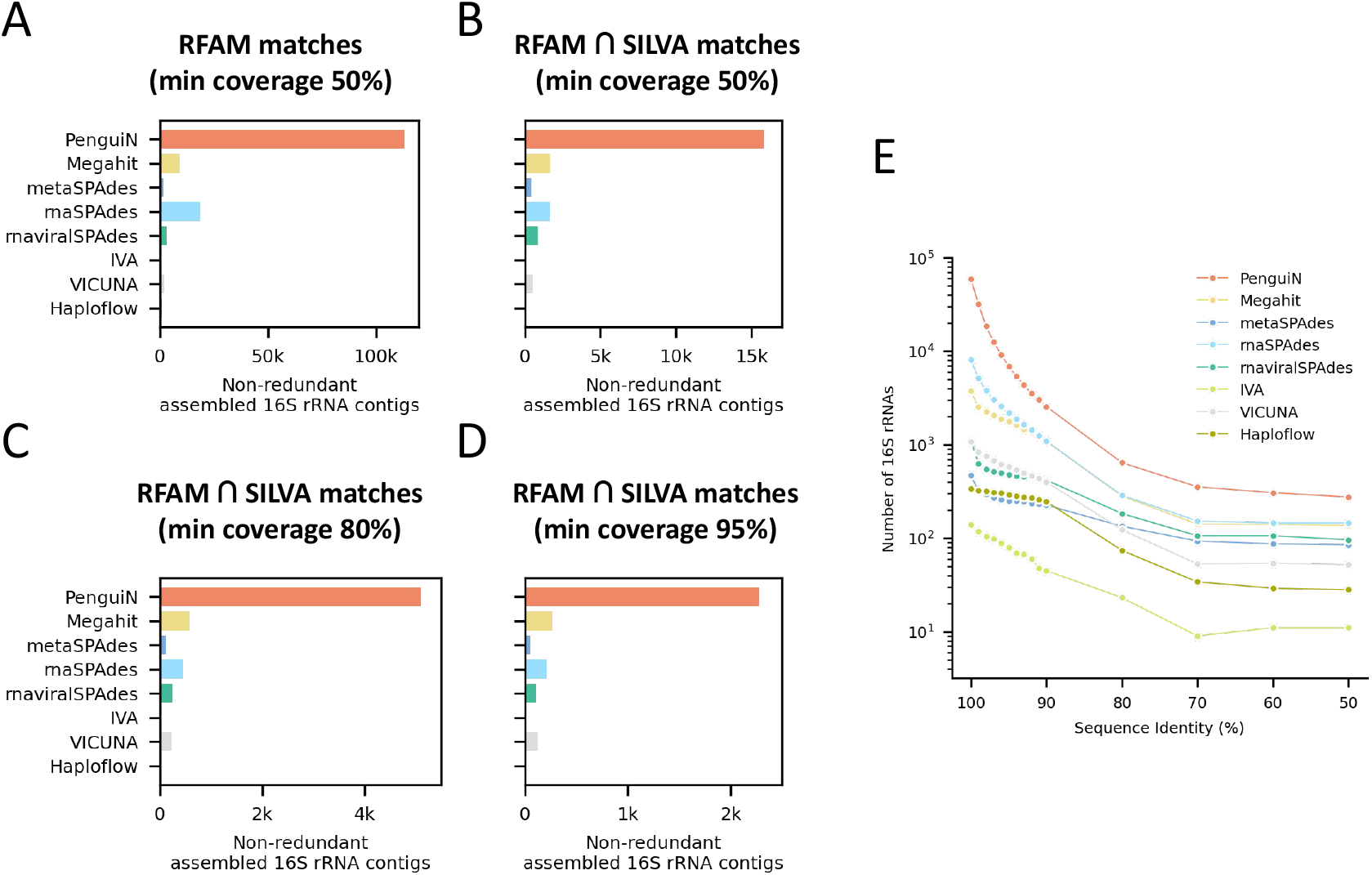
Number of assembled 16S rRNA contigs matching RFAM and SILVA entries. (**A**) Non-redundant, assembled 16S rRNAs matching the RFAM 16S HMM model with at least 50% of coverage. (**B-D**) Non-redundant, assembled 16S rRNAs additionally validated by matching a 16S rRNA sequence in the SILVA database with at least 99% sequence identity and covering at least 50% (B), 80% (C) and 95% (D) of the SILVA reference sequence. (**E**) Number of representative, assembled 16S rRNA sequences matching the RFAM 16S HMM model with >80% of coverage after removing redundancy by clustering at different levels of sequence identity.

To ensure that the assembled contigs represent correct, non-chimeric 16S rRNA sequences, we validated the assembled contigs by MMseqs2 searches [49] through the SILVA database, a comprehensive database containing 510 495 full-length, quality-checked 16S/18S rRNA sequences [54]. We counted how many contigs in each assembly could be mapped to a SILVA 16S rRNA sequence with at least 99% sequence identity and a coverage of the SILVA sequence of at least 50%, 80% and 95% (Fig. **7**B,C,D). The analysis using only these validated sequences reveal the same pattern: At each coverage threshold, PenguiN assembles around 6 to 8 times more 16S rRNA genes than the next best tool. For example, PenguiN assembled 2284 16S rRNAs covering ≥ 95% of the full length reference in SILVA, while the next best tool, Megahit, assembled 269, or 8.5 times less (Fig. **7**D).

The diversity of the 16S rRNA genes reconstructed by PenguiN was several times higher over a large range of sequence identities (Fig. **7**E). For example, filtering representative sequences using 97% maximum pairwise identity, PenguiN’s assembly contained 12 616 16S rRNA genes while the next best tool, rnaSPAdes, assembled only 3068. At 99% clustering threshold, corresponding roughly to the species level [55], PenguiN reconstructed *∼* 6 times more species-level 16S rRNAs than rnaSPAdes (32 169 versus 5132). Remarkably, PenguiN assembled at least twice more 16S rRNA sequences than the next best tool even down to 50% clustering threshold.

### Run time and memory usage

Previous overlap-based approaches are computationally expensive due to the quadratic complexity of all-versus-all alignments and were therefore superseded by de Bruijn graph assembler in metage-nomics. For each assembler and dataset, we measured the run time and peak memory usage for the assembly processes (see Table 1 and Table 2) on a 16-core server with two Intel Xeon E5-2640v3 CPUs and 128 GB memory. Assemblies were performed individually for each sample with multithreading options set to utilize all 16 cores. Runtime was limited to 10 days per sample.

**Table 1:** Comparison of runtimes (wall times) for all datasets used in the assembly benchmark. All tools were run with 16 threads, except for Haploflow, which did not support multi-threading. Values are given in [hh]:mm:ss. ‘-’: assemblers did not produce any results. Values marked with * are lower bounds because the tools did not finish on all samples.

| Dataset / Tool | HRV in silico mixture | HIV1 in silico mixture |  |  | Metatranscriptomic dataset |
| --- | --- | --- | --- | --- | --- |
|  |  | 1x | 10x | 100x |  |
| PenguiN | 00:08 | 00:18 | 00:03:34 | 00:25:40 | 89:17:51 |
| Megahit | 00:03 | 00:33 | 00:03:14 | 00:08:36 | 28:39:21 |
| metaSPAdes | 00:19 | 02:14 | 01:02:34 | 02:12:02 | 668:47:30 |
| metaviralSPAdes | 00:23 | 01:02 | 00:04:03 | 00:13:39 | - |
| rnaSPAdes | 00:08 | 00:27 | 00:02:14 | 00:08:06 | 34:28:50 |
| rnaviralSPAdes | 00:11 | 00:37 | 00:03:32 | 00:10:08 | 31:31:44 |
| SAVAGE | 01:41 | 25:27 | - | - | - |
| IVA | 03:04 | 04:31 | 00:51:31 | 29:17:30 | - |
| VICUNA | 00:02 | 00:33 | 03:28:33 | 29:31:19 | *4989:38:20 |
| Haploflow | 00:02 | 00:47 | 00:03:41 | - | *568:57:30 |

**Table 2:** Comparison of Max RAM usage for all datasets used in the assembly benchmark. Values are given in GB. ‘-’: the assembler did not produce any results. *Haploflow had one sample in the metatranscriptomic dataset where the 128 GB were not sufficient for the assembly, resulting in an “out-of-memory error”.

| Dataset / Tool | HRV in silico mixture | HIV1 in silico mixture |  |  | Metatranscriptomic dataset |
| --- | --- | --- | --- | --- | --- |
|  |  | 1x | 10x | 100x |  |
| PenguiN | 0.10 | 0.56 | 2.69 | 15.08 | 91.01 |
| Megahit | 0.25 | 0.27 | 0.29 | 1.81 | 10.96 |
| metaSPAdes | 5.11 | 5.15 | 22.16 | 46.97 | 116.66 |
| metaviralSPAdes | 5.11 | 5.26 | 5.67 | 11.59 | - |
| rnaSPAdes | 5.11 | 5.21 | 5.49 | 11.55 | 24.58 |
| rnaviralSPAdes | 5.11 | 5.21 | 5.47 | 11.55 | 17.64 |
| SAVAGE | 1.28 | 10.94 | - | - | - |
| IVA | 1.25 | 1.24 | 1.24 | 1.25 | - |
| VICUNA | 0.06 | 0.35 | 3.23 | 31.03 | 116.38 |
| Haploflow | 0.01 | 1.06 | 1.79 | - | *119.00 |

All assembly tools were able to assemble the small HRV dataset in very short time. On the HIV1 dataset, we see the trend that the de Bruijn graph assemblers are much faster than all overlap-based tools except PenguiN. As expected, these overlap-based approaches showed severe limitations on the large and complex metatranscriptomic samples. SAVAGE and IVA did not produce any results within 10 days for these datasets and VICUNA could only process 33 of the 82 samples within 10 days. PenguiN was the only overlap-based assembler that could process this dataset completely. Overall, the de Bruijn graph assemblers Megahit and rna(viral)SPAdes were the fastest assemblers. PenguiN was 3 times slower, but 5-7 times faster than metaSPAdes and much faster than VICUNA (> 55 times).

PenguiN’s memory usage was larger than that of Megahit and rna(viral)SPAdes, but still not limiting on our large and complex datasets, and smaller than that of metaSPAdes, VICUNA and Haploflow.

## 3 Discussion and Conclusion

PenguiN is the first overlap-based tool for strain-resolved viral genome assembly from metagenomic data. We compared it to nine state-of-the-art metagenomic and viral genome assemblers. On a simulated dataset with reads from 2550 HIV-1 genomes, PenguiN assembled a fraction of the HIV-1 genomes – matched at 99% sequence identity – that was many times higher and with much lower fragmentation than the next best assembler rnaSPAdes. Further, PenguiN assembled around three times more complete ssRNA phage genomes from 82 samples from activated sludge and aquatic environments. PenguiN also assembled around six times more full-length, bona-fide 16S rRNA sequences from the 82 metagenomic samples than the next best tool, rnaSPAdes.

Despite PenguiN’s high sensitivity, its precision (the fraction of assembled contigs that could be matched to one of the reference genomes) was similar to the state-of-the-art tools. These results also suggests that the fraction of chimeric contigs is similar to the other assemblers because chimeric assemblies should result in lower precision. Indeed, on the metagenomic ssRNA phage genome data, none of the 1398 complete phage genomes assembled by PenguiN contained genes with discordant phylogenies, indicating that no chimeric genomes were assembled. Furthermore, comparison of the number of assembled 16S rRNA contigs before and after validation using mapping to SILVA reference sequences (Fig. **7**A vs. B) demonstrates that the fraction of contigs that could be validated – and therefore are not chimeric – is very similar for PenguiN (14%) and Megahit (19%) and higher than for rnaSPAdes (9%). We conclude that, despite or perhaps rather because of its greedy extension approach, PenguiN produces a similar fraction of chimeric contigs as its best competitors.

When dealing with viral metagenomic data, the choice is between de Bruijn graph based metagenomic assemblers and virus-specific assemblers for low-complexity samples that mostly use overlap graphs. Our benchmarking results agree with results described in other studies [32, 56]: Due to the limitation of sequence context to the *k*-mer length, de Bruijn graph-based assemblers cannot phase highly similar genomes, while overlap-based assemblers often cannot handle large and complex metagenomic samples.

PenguiN’s use of the Linclust algorithm allows it to overcome the quadratic runtime complexity that plagued earlier overlap-based assemblers. Our results show that PenguiN can process large and complex datasets in reasonable time, albeit thrice slower than the fastest de Bruijn graph assemblers. In contrast, the other overlap assemblers we tested needed either inordinate amounts of time or failed on the dataset of 82 metagenomics samples.

Due to utilizing overlap alignments, PenguiN can make use of a much longer sequence context, the length of merged read pairs, in contrast to de Bruijn graph assemblers, whose sequence context length it limited to the *k*-mer length. Due to its Bayesian model to select the best sequences, PenguiN can exploit this longer context efficiently. The trend to increasing short read lengths – Illumina’s current high-throughput sequencers NovaSeq 1000/2000 and 6000 can generate paired reads up to 2×300 and 2×250 base pairs, respectively – will further increase the length of PenguiN’s sequence context and thus further improve its strain resolution compared to our benchmarks based on 2 × 150 paired reads.

In the same vein, PenguiN should profit a lot from long-read sequencing platforms such as Oxford Nanopore. The important caveat is that, due to its gapless alignment procedure during extension, it is only suitable for reads with indel rates similar to those for Illumina machines. Two recent approaches reach sufficiently low indel errors: In read polishing, Illumina short reads are mapped to long reads to correct errors [57], and Nanopore duplex sequencing reads both strands of a DNA several times [58].

Due to PenguiN’s Bayesian selection and contig extension strategy, strain resolution is optimal when the coverage for a strain genome is 10 or more, as in that case a read from the correct strain with good overlap to the contig is likely to be present in the dataset. When the coverage drops much below 10, the greedy extension strategy automatically falls back to reconstructing consensus sequences like other metagenomic assemblers do [44]. Our results show that, at low, medium and high read coverage, PenguiN produces similar amounts of assembly errors as the state of the art.

PenguiN has two main drawbacks. First, as shown in Fig. 3b, its assemblies are slightly redundant, to a similar degree as the next best assembler in our benchmarks, rnaSPAdes. Second, and most importantly, PenguiN does not stop with its greedy extension when extensions are ambiguous. Therefore, it performs poorly when *identical* repeats or 100% conserved regions longer than the paired-read length are present in the metagenomes, since in this case PenguiN will randomly connect sequences upstream with sequences downstream. This seems to be no limitation for viral assembly, but in preliminary experiments, this limitation led to suboptimal assemblies of some prokaryotic genomes. We are planning to address this limitation in the future by terminating the extension process when PenguiN detects a bifurcation. This might be done by combining PenguiN’s linear-time overlap determination with an overlap graph-based approach. As a future improvement for assembling strand-specific metatranscriptomic libraries, an option could be added to perform strand-specific assembly in which reverse complements of reads during extension are not allowed.

In conclusion, PenguiN is the first overlap-based assembler for viral genome and 16S rRNA assembly from large and complex metagenomic datasets. Due to analysing all overlap alignments and a Bayesian strategy to select optimal extensions, it is able to assemble several fold more strain-resolved genomes than the state of the art. The last decade has brought fascinating discoveries of a huge and often unexpected diversity of viruses in many different environments [6, 7, 8, 9, 10, 11, 12, 13, 14, 15, 16, 17, 18], and a large part of these successes were driven by progress in viral bioinformatics and computational tool development [59, 60, 61, 22]. We hope that PenguiN will make an important contribution to studying the roles of viruses in earth’s microbiomes and to uncovering the molecular machineries evolved by these most diverse and inventive of all biological entities.

## 4 Methods

### The protein-guided nucleotide assembly approach

PenguiN proceeds in two stages (Fig. **1**): In the protein-guided stage, it assembles six-frame translated protein sequences and co-assembles the corresponding nucleotide sequences. The resulting coding sequence-containing contigs are added to the original nucleotide reads and further assembled in the second stage. Each of the two stages consists of iterating the following two steps: (1) Find the overlaps among the working set of sequences that satisfy the specified criteria for extensions (maximum E-value, minimum sequence identity, and number of aligned residues), and (2) find the best upstream and downstream extensions to merge, and add the merged sequences into the working set. The two stages are now described in more detail (Additional File 1: Fig. S5).

In stage I, paired-end reads are first merged, then six-frame translation is performed by traversing each read linearly and using a standard codon table. Subsequently, potential ORFs with ≥ 45 codons are extracted from the translated sequences. Additionally, we extract ORFs with at least 20 codons starting with a putative ATG start codon, i.e. the first ATG codon after a stop codon in the same frame. Then five iterations of overlap identification and extension are performed. First, protein overlaps are found in a time proportional to the number of sequences using the following Linclust algorithm [45]: A fixed number of *k*-mers (default: 60) are extracted from each sequence and stored in a reverse index table via a hash function value. This table is used to group sequences that share an identical *k*-mer substring (*k* = 14). The longest sequence per group, called center sequence, is selected, and groups with the same center sequence are merged together. Each center sequence is then aligned to each of the member sequences in its group. Because group member sequences are not aligned with each other, the algorithm achieves linear rather than quadratic time complexity. As in the protein-level assembler Plass [28], the alignments that satisfy the specified criteria of maximum E-value, minimum amino acid sequence identity, and minimum nucleotide sequence identity, are further considered as sequence extensions. Since we want to minimize the risk of chimeric proteins that would lead to falsely connected nucleotide sequences, we chose strict default filter criteria of E-value < 10^−5^, amino acid identity > 97%, and a nucleotide identity of > 99%. We select the best upstream extensions and the best downstream extensions according to a Bayesian posterior probability (see below). Each time two protein sequences are merged for extension, the corresponding nucleotide sequences are also extended in parallel. This stage ends after five iterations with pre-assembled contigs containing coding sequences. As in Plass, sequences translated in wrong frames tend to result in short protein contigs due to frequent stop codons. Their presence does not help, but also does not hurt, as the resulting short contigs are eliminated during the length and subsequent redundancy filtering at the end.

In stage II, PenguiN links these pre-assembled contigs across intergenic regions. The coding sequence contigs are added to the original reads, and five iterations using the same greedy assembly procedure as in stage I are applied to the nucleotide sequences. To keep the specificity sufficiently high, a nucleotide *k*-mer size *k* = 22 is used in this stage. PenguiN again searches for overlaps in linear time and decides on the best sequence extensions based on the full alignments of the overlapping sequences. Again, the center sequences to be extended are the longest sequences in each group of sequences sharing a *k*-mer seed. In this way, we primarily extend contigs with other contigs and with non-coding reads, and, if there are no more pre-assembled contigs within the group, we also extend non-coding reads with non-coding reads. Since sequences and their reverse complements code for the same double-stranded DNA, we identify each *k*-mer with its reverse complement. We replace group member sequences with their reverse complement if their *k*-mer is the reverse complement of the matching *k*-mer in the center sequence. To maintain high sensitivity for identifying the overlaps, we scale the number of extracted seeds with the length of the sequences, by default we extract 0.1× the number of residues in the sequence. In addition, we check for circular contigs after each iteration in stage II, because plasmids and many viral genomes are circular (see below). Since we cannot distinguish between circular genomes and perfect repeats, PenguiN will also stop extending contigs framed by two perfect repeats. Only the non-circular contigs are added to the working set to be further extended, while the circular ones are excluded from subsequent iterations and added directly to the output contigs.

For the final output, circular contigs collected over all iterations and linear contigs from the last iteration are combined. Since PenguiN is able to use reads more than once, the set of final contigs may be quite redundant. Therefore, in the last step, we perform a clustering of all final contigs using an extended version of Linclust [45] that takes circular contigs into account (option –wrapped-scoring), using a default maximum sequence identity of 99% and minimum coverage of 99% to obtain strain-resolved contigs. The longest sequence in each cluster is returned as result.

### Choosing extensions

PenguiN uses a Bayesian model to decide with which of several alternative choices of overlapping sequences to extend a center sequence. When overlap alignments differ in length, this model improves upon the method used in Plass [28], which simply picks the extension with the highest sequence identity. We can calculate the probability that the genome from which one extending sequence was sampled has a higher sequence similarity with the genome from which the center sequence was sampled than the other extending sequence.

Suppose we have a possible extension with *L* overlapping, aligned nucleotides, of which *m* are matches and *L* − *m* are mismatches. The probability to observe *m* matches among *L* aligned residues is given by a binomial probability,

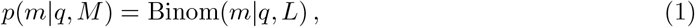

where *q* is the unobserved, unknown average nucleotide identity (ANI) between the strains from which the two sequences originate. As prior for *q* we use a Beta distribution *p*(*q*) = Beta(*q*|*a, b*) with pseudocounts *a* and *b*, since it is the conjugate prior to the Binomial distribution and the simplest smooth distribution on the interval [0, 1] that can model any possible mean and variance. Using Bayes’ theorem, we can calculate the posterior probability distribution of the ANI *q* (see Additional File 1: Supp. Note 1),

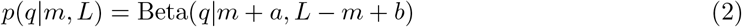

Now suppose we have to decide which of two possible extending sequences to choose, one with *m*_1_ matches out of *L*_1_ aligned, overlapping residues and one with *m*_2_ matches out of *L*_2_ aligned residues. The probability that the ANI *q*_1_ between the contig and the first extending sequence is lower than the ANI *q*_2_ between the contig and the second extending sequence is

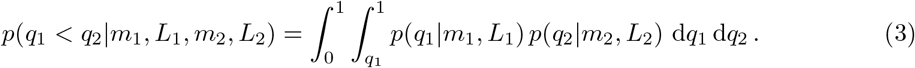

The Supplemental Material shows how to analytically solve this integral and compute it efficiently.

To decide if we want to preferably extend with an alignment (*m*_1_, *L*_1_) rather than (*m*_2_, *L*_2_), we proceed as follows: If *p*(*q*_1_ > *q*_2_|*m*_1_, *L*_1_, *m*_2_, *L*_2_) > 0.55 the first sequence is preferred for the extension, if *p*(*q*_1_ > *q*_2_|*m*_1_, *L*_1_, *m*_2_, *L*_2_) < 0.45 the second one is preferred. If the probability falls into the range 0.45 to 0.55 we consider it as “inconclusive”, and prefer the extension that offers the longer extension instead. The center sequence is extended by the member sequence of its group that is preferred over all other possible extensions. For the pseudocount parameters we choose *a* = 1, *b* = 1.

### Cycle detection

As circular and long terminal repeats are common among viruses and would lead to over-extensions in the greedy iterative assembly procedure, we introduced a step to detect contigs containing such structures. Therefore, we search for contigs with (near-)identical terminal repeats on both sides and exclude them from the next iterations. Instead, we mark them as circular and collect them over all iterations - marked by a flag in their header entry. To identify circular sequences while minimizing the runtime, we use a procedure that approximates the alignment of the sequence to itself using *k*-mer matches. We look for an overrepresentation of *k*-mer matches within diagonal bands and mark the contig as circular if a hit rate threshold of 0.24 is exceeded. This threshold was calculated using 5994 viruses from the RefSeq [62] database.

### Simulated datasets / Assembly benchmark

For benchmarking experiments, we generated synthetic read datasets. To make the distribution of the coding sequences as realistic as possible, we generated the reads from real viral genomes from the NCBI database [63]. For the first set (HRV in silico mixture), we downloaded three human rhinovirus strains (Accession No.: MF973193.1, MF973194.1, MN749156.1) with pairwise ANI values ranging from 92 to 95.5%. We mixed them in a proportion of 4:2:1 and simulated 2 × 150 bp overlapping paired-end reads (insert size range 220-280 bp) using randomreads.sh from the BBmap software suite (version 38.71) [64]. The simulated coverage was set to 50.

For the HIV datasets, we aimed to construct benchmark sets with a high degree of natural variation. We downloaded 2550 HIV-1 genomes from the NCBI database (accessed 07/2019, search string: (“Human immunodeficiency virus 1”[Organism] OR hiv1[All Fields]) AND complete genome[All Fields]). Comparing their ANI values with fastANI showed values mostly in the range from 85-95% (see Fig. **3**A). We then used randomreads.sh to generate three synthetic read datasets from this set using varying coverage values (1, 10, 100). Next, we also introduced sequencing errors with an error rate of 0.001%, 0.01%, 0.1% for the 10× coverage set to check if the assembly quality is influenced by the presence of sequencing errors.

The HIV1 genome is approximately 9.7 kbp and is flanked at both ends by long terminal repeats (LTRs) that can interact, leading to circularized forms containing one or two copies of the viral LTR (1-LTR circles, 2-LTR circles) [47, 48]. All downloaded 2550 HIV-1 genomes were reported as “complete genome” in the NCBI database, but their LTRs were reported inconsistently (without, with one LTR or two LTRs, etc.). Because of that, it was not clear what result an assembler should ideally produce (genome with one LTR or two LTRs, etc.). Therefore, we consistently circularized all genomes by removing one of the repetitive regions, if two were present, and doubling the sequences, before simulating the reads. This results in reference genomes to which all linear representations of the complete genome can be mapped continuously, regardless of the starting position of the assembled contig.

We assembled all synthetic paired-end read datasets using PenguiN, Megahit, metaSPAdes, metaviral-SPAdes, rnaSPAdes, rnaviralSPAdes, SAVAGE, IVA, VICUNA, and Haploflow.

### Evaluation on simulated datasets

For the evaluation of the assemblies, only contigs > 1 kb were considered. Evaluations of assemblies were performed with MetaQUAST [46], which provides commonly used statistics. We used MetaQUAST with the `--unique-maping` flag and default parameters otherwise as it is commonly used in metagenomic assembly benchmark studies [65, 32, 66]. In addition, we computed the per-base sensitivity and per-base precision values for the HIV datasets as we previously defined [28]. We searched with the set of reference genomes through the assembled contigs using MMseqs2 [49] with options -a -s 5.7 `--max-seqs 500000 --min-seq-id 0.89 --strand 2--search-type 3 --max-seq-len 1000000` and subsequently filtered the resulting alignments with a minimum sequence identity threshold between 90% and 99%. Sensitivity was then calculated as the fraction of reference sequences that can be covered by contigs with a sequence identity of at least this threshold (total count of aligned nucleotides divided by the total length of the reference genomes). We considered two ways - either all alignments or only the longest alignment for each reference genome. Precision was calculated by searching with the contigs through the set of reference genomes, using MMseqs2 with options `-a -s 5.7 --max-seqs 5000 --min-ungapped-score 100 -a --min-seq-id 0.89 --strand 2 --search-type 3 --max-seq-len 10000000`. For each contig, only the longest alignment was considered. Precision was then defined as the fraction of contigs that are aligned to the reference genomes (total count of aligned nucleotides divided by total length of assembled contigs).

### Metatranscriptomic datasets

For the benchmark test on real sequencing data, we considered 82 metatranscriptomic samples from activated sludge and aquatic environments previously used in [50]. We downloaded the samples from the NCBI Sequence Read Archive (SRA) database and pre-processed them analogously to the Callanan et al. study. We removed Illumina adapters using Cutadapt (version 3.3) [67] and performed quality trimming using Trimmomatic (version 0.39) [68]. We discard all reads shorter than 100 bp. Assembly was then performed per sample using PenguiN, Megahit, metaSPAdes, metaviralSPAdes, rnaSPAdes, rnaviralSPAdes, SAVAGE, IVA, VICUNA, and Haploflow. The minimum contig length was set to 500 and the maximum runtime per sample was set to 10 days. We found that SAVAGE and IVA were unable to process the samples, so they were excluded from the results.

### Detecting ssRNA phage sequences in the assemblies

To detect and classify ssRNA phage contigs in the assemblies of the metatranscriptomic samples, we used the HMM based pipeline described in Callanan et al. [50] after the assembly. We predicted proteins with Prodigal (version 2.6.3) [69] and searched against the sequence profiles of the HMM 5-MC model generated by [50] using hmmscan from the HMMER package version 3.2.1 at http://hmmer.org/. Clustering was performed using MMseqs2. The sequence identity cutoff for clustering was first chosen as 100% in accordance to Callanan et al. However, for comparison between assemblers, we chose a cluster identity threshold of 99% to avoid overestimating sequences close to duplicates. All sequences longer than 750 bp were classified depending on the number of protein hits. We classified the contigs using the same terms as Callanan et al.: contigs encoding at least two protein hits (“partial genomes”), contigs encoding three protein hits (“near-complete genomes”), and contigs encoding three protein hits without proteins that are prematurely terminated by the edge of a contig (“complete genomes”).

### Analysis of ssRNA phage sequences

Due to the lack of ground truth for real data, we could not calculate precision or sensitivity values, instead we assessed the assembly quality by analysing the complete ssRNA phage sequences in more detail. We calculated the fraction of PenguiN’s assembled phage sequences which are covered by alignments to phage sequences in the other assemblies at a minimum identity cutoff of 99%. Aligning was performed using MMseq2 search [49] with options `--max-seqs 500000 -a --min-seq-id 0.99 --strand 2 --search-type 3 --min-aln-len 300 --max-seq-len 1000000`. For two assemblies *A* and *B* we counted a contig from *A* to lie in *A ∩ B* if it has an alignment with a contig from *B* with ≥ 99% sequence identity and ≥ 99% coverage. To analyse the recovered diversity of the different assemblies we extracted from all complete phages the amino acid sequences that had a hit to the RdRp HMM model, and clustered them using MMseqs2 cluster [49] at sequence identity cutoffs of 50-100% (with step = 10 until 90% and step = 1 between 90 and 100%). To check the phylogenetic consistency we considered the subgroups of the three core genes that were observed to co-occur only in specific patterns [50]. We checked the order and co-occurrence of the core genes for deviations from these patterns using the list of predicted proteins provided by Prodigal and their respective HMM scores.

### Detecting 16S rRNA gene sequences in the assemblies

To detect 16S rRNA genes in the assemblies of the 82 metatranscriptomic samples, we used Barrnap v.0.9 (https://github.com/tseemann/barrnap). We classified predictions as being aligned to at least 50% of the 16S HMM model from the RFAM database v.11.0 [70]. To reduce redundancy, predicted 16S rRNA sequences were clustered using MMseqs2 with a sequence identity cutoff of 99%. To validate the assembled 16S rRNA contigs and determine the number of full-length 16S rRNAs recovered, we searched the 16S contigs (non-redundant) against the SILVA database v.138.1 [54]. The search was performed using MMseqs2 with a minimum sequence identity cutoff of 99% and coverage threshold of 50%, 80% and 95%. To estimate the diversity of the 16S rRNA contigs, we clustered the sequences predicted by Barrnap using MMseqs2 with sequence identity cutoffs of 50-90% (with step = 10) and 91-99% (with step = 1). We then counted the number of cluster representatives having more than 80% of their sequence aligned to the Barrnap 16S HMM model.

### Software

The respective software versions of the assembly tools used in the benchmark are: PenguiN (GitHub commit 7571d37), Megahit (v1.2.9), metaSPAdes/rnaSPAdes/metaviralSPAdes/rnaviralSPAdes (v3.15.2), SAVAGE (v0.4.2), IVA (v1.0.8), VICUNA (v1.3), Haploflow (v0.1).

Each assembler was run with default parameters, except for the minimum contig length and the CPU thread parameter setting, which was uniformly set to utilize all 16 available cores of the machine, if possible. The minimum contig length was set to 500 bp or 1000 bp depending on the dataset for all assemblers that provide such a filter option, or otherwise by filtering the final assembly file. As input, we used paired-end reads in two separate files or concatenated them if necessary.

### Data availability

All scripts and benchmark data including command-line parameters necessary to reproduce the benchmark and analysis results presented are available at https://github.com/AnnSeidel/penguin-analysis.

### Code availability

We integrated PenguiN into our Plass software. Source code and binaries are freely available under GPL-3.0 licence from https://github.com/soedinglab/plass. The latest released version is 5-cf8933. The source code used for benchmarking in this article can be accessed as git revision commit 7571d37. PenguiN is written in C++ and requires a 64-bit Linux/macOS system with at least SSE2 instructions

## 5 Declarations

## Acknowledgements

We thank Louis Kraft and Jinlong Ru for early testing of the software and giving feedback for parameter settings, and Bas Dutilh for feedback on the manuscript. JS acknowledges support by the ERC’s Horizon 2020 Framework Programme [‘Virus-X’, project no. 685778] and the BMBF CompLifeSci project horizontal4meta. MS acknowledges support from the National Research Foundation of Korea (grants 2019R1A6A1A10073437, 2020M3A9G7103933, 2021R1C1C102065, and 2021M3A9I4021220), the Samsung DS Research Fund, and the Creative-Pioneering Researchers Program through Seoul National University.

## Author contributions

JS and MS conceived and supervised the research. AJ and MS developed the software. JS and EJ developed the Bayesian model for extension selection. AJ performed the virus data benchmarks. AJ and FAJ tested the software and analysed results. AK performed the 16S rRNA analysis. AJ and JS wrote the manuscript.

## Ethics approval and consent to participate

Not applicable.

## Consent for publication

Not applicable

## Supplementary Information

### Supplementary Figures

**Figure S1:**
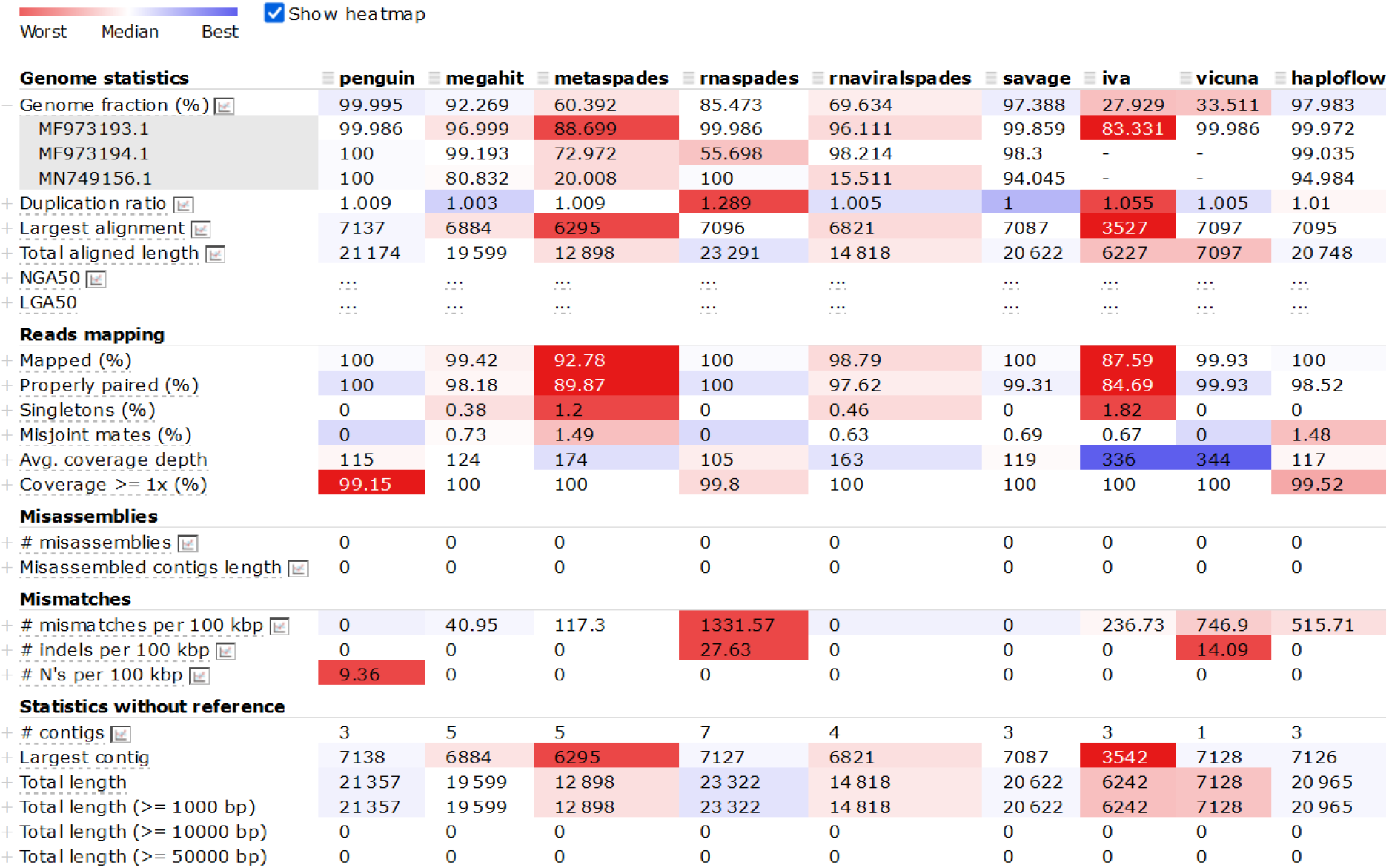
MetaQUAST assessment of assembly quality on an in-silico mixture of three HRV genomes.

**Figure S2:**
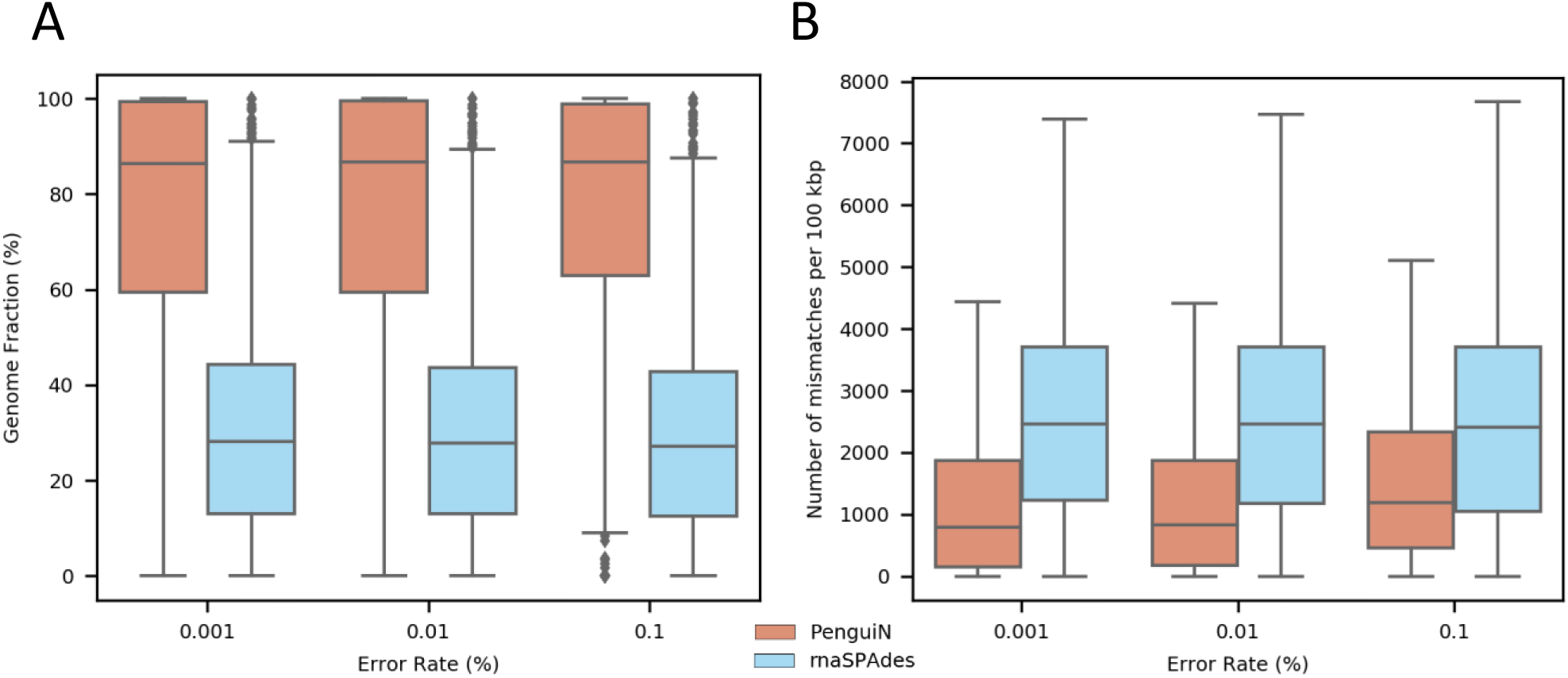
Influence of sequencing errors on the assembly performance for the HIV1 10× coverage dataset evaluated by MetaQUAST. For better visualization we did not plot the outliers in subfigure B.

**Figure S3:**
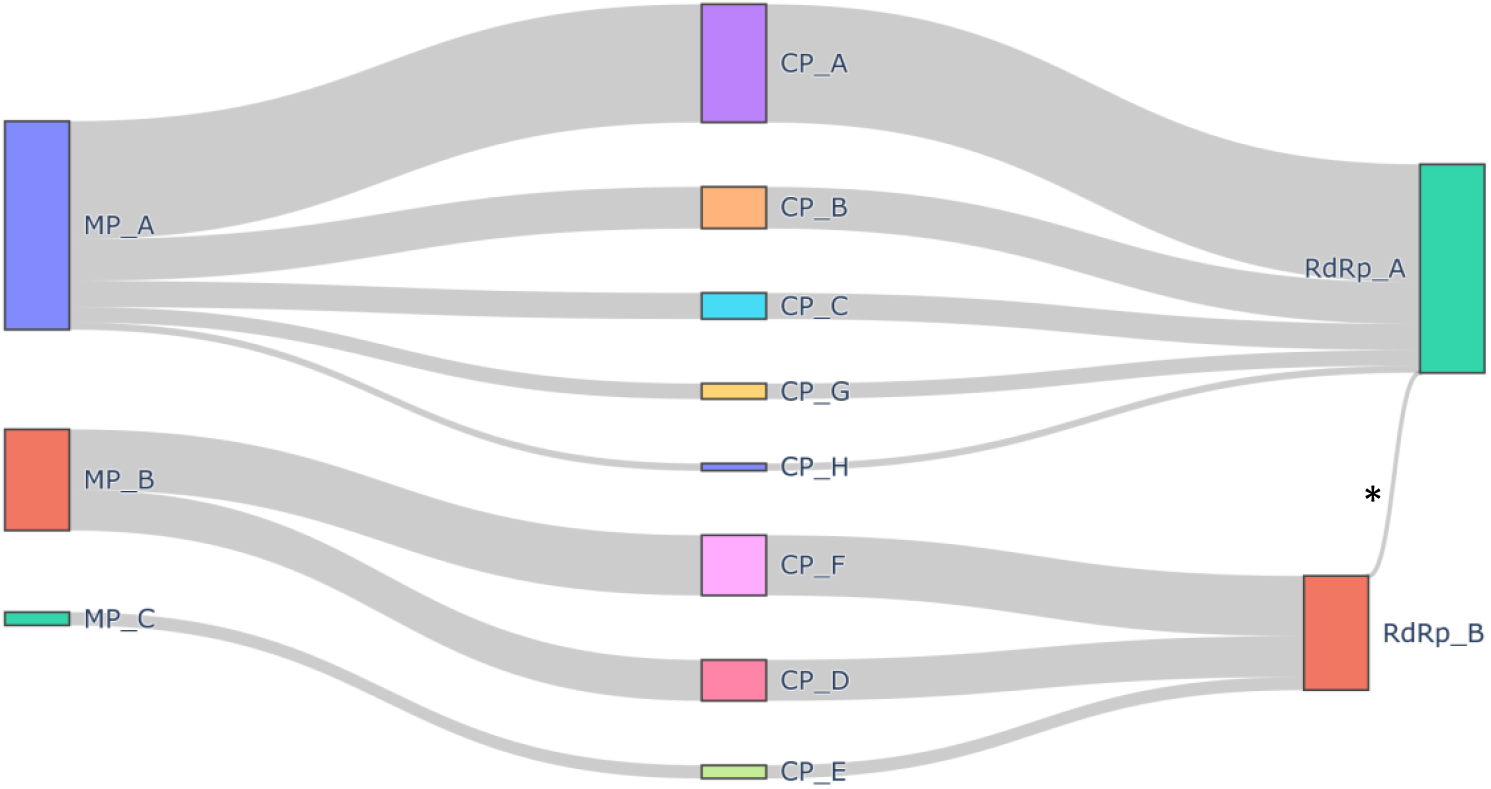
Co-ooccurence profiles of the core proteins in complete ssRNA phage genomes assembled with PenguiN. The HMM profiles released by [1] contained 3 MP, 8 CP and 2 RdRp protein clusters. Of the 48 theoretical combinations of these proteins on a complete genome, only 8 were observed in the PenguiN assembly. This is in accordance to what was seen in the Callanan et al. study. This indicates that the assembled sequences have proteins with concordant phylogenies, in contrast to what we would expect from missassemblies or chimeric sequnces. There was only one deviation marked by an asterisks (*) where a contig (contig: SRR7976310 76852) contained an additional RdRp A following the RdRp B. However, the protein classified as RdRp A (hmmscan score 469.6) in this contig was also very similar to the RdRp cluster B (hmmscan score 240.0)

**Figure S4:**
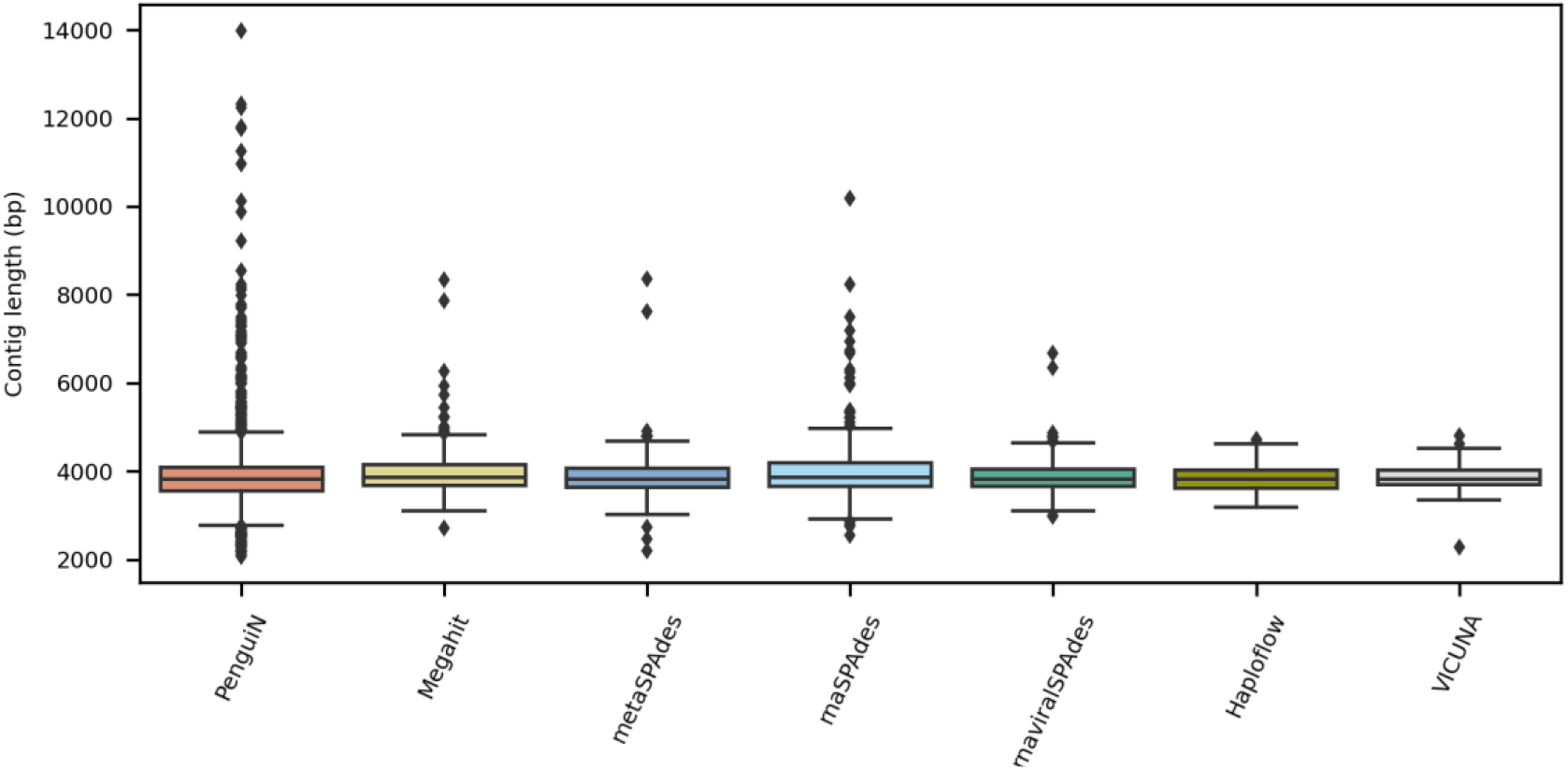
Length distribution of the complete ssRNA phage genomes identified from the different assemblies.

**Figure S5:**
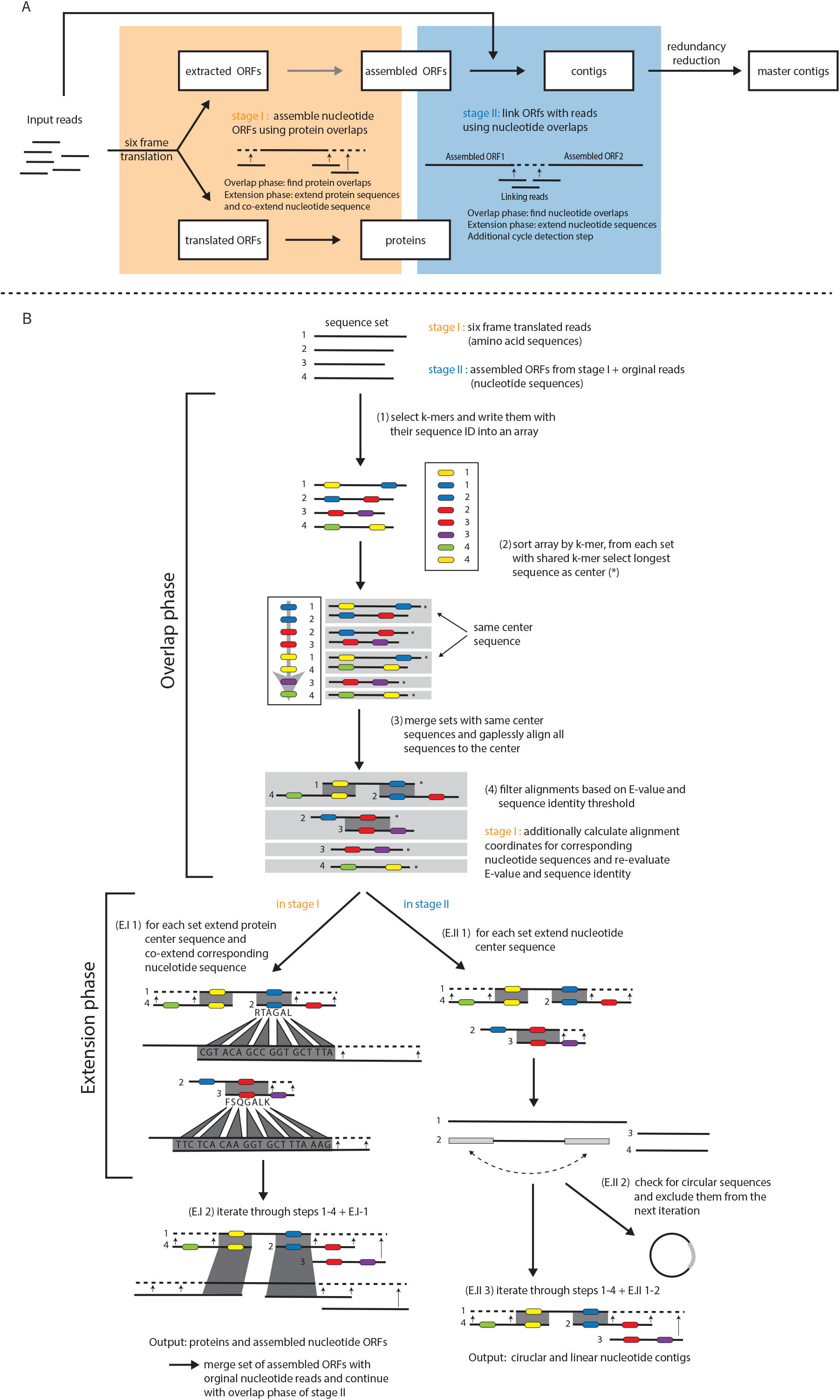
The stages of PenguiN’s assembly in more detail.

### Supplementary Notes

**Supplementary Note 1: Greedy extension strategy based on a Bayesian model**

In the extension step, each center sequence is extended by concatenating the non-overlapping residues of the member sequences on each side (“hang-offs”), either on protein and nucleotide level simultaneously (stage 1), or on nucleotide level only (stage 2).

Even though only the member sequences with alignments to the center sequence satisfying the sequence identity threshold and the E-value criterion are considered, multiple extensions might be possible on each side of a center sequence. In the Plass [2] algorithm, the list of alignments is sorted in order of descending overlap sequence identity and the best left and right extensions are chosen. However, the length of the compared overlaps can vary greatly, which is particularly seen during the nucleotide assembly or in later iterations of the guided assembly.

For example, an overlap with length 1000 and sequence identity 98.9% might be more trustworthy than an overlap of length 100 and sequence identity 99.0% as a longer overlap also yields a statistically more significant estimation of the actual similarity between the full sequences. We used a Bayesian formulation of the problem, which can be solved analytically.

Given a query sequence (center sequence) and a target sequence (member sequence) with a mean fraction of identical residues *q*, and an alignment of length *L* between them, out of which *m* are matches and *L* − *m* are mismatches, the probability distribution for the number of matches in the alignment is a binomial distribution

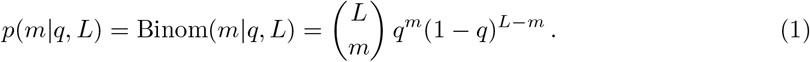

The *q* are hidden variables while *L* and *m* are the observed variables. Given the latter, the probability distribution of the former can be deduced using *Bayes’ theorem*,

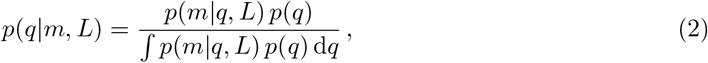

where *p*(*q*) is the prior probability for the fraction of matches of the alignment. We model *p*(*q*) as a beta distribution,

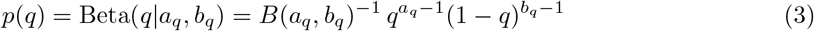

With

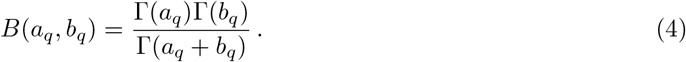

We therefore obtain for the hidden *q*

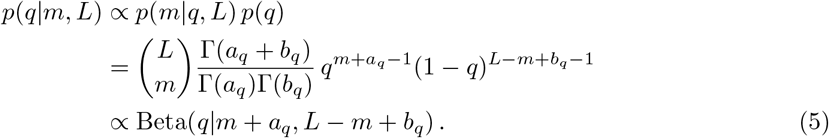

The proportionality constant between the left and right-hand sides must be 1 since both are normalized probability distributions.

Given two possible extensions for the same contig, described by the alignments (*m*_1_, *L*_1_) and (*m*_2_, *L*_2_), the probability that the first target sequence has lower average nucleotide identity (ANI) to the query contig than the second, is then given by

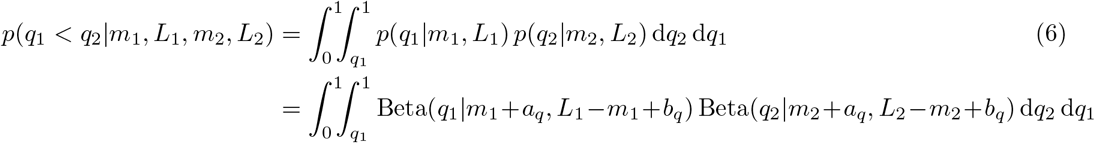

using formula (5). The probability *p*(*r*|**m, L**) that *r* is the best matching sequence extending the contig is

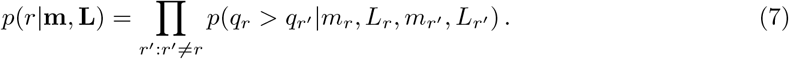

To evaluate the probabilities *p*(*q*_1_ < *q*_2_|*m*_1_, *L*_1_, *m*_2_, *L*_2_), we need to compute the integral

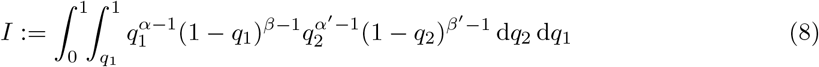

for

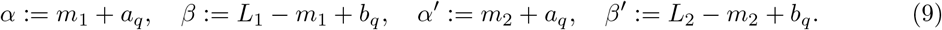

With a change of variables from *q*_2_ to *t, q*_2_ = *t* + (1 − *t*)*q*_1_; *t ∈* [0, 1] with d*q*_2_ = (1 − *q*_1_) d*t*, the integral can be rewritten with fixed boundaries,

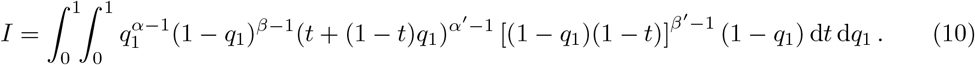

Expanding

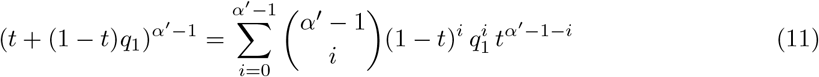

yields

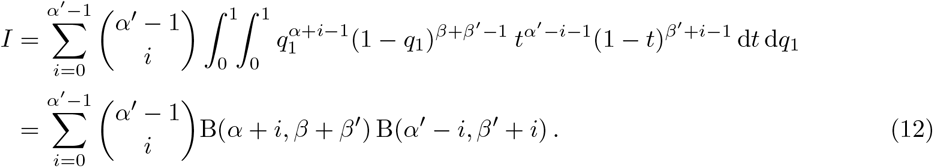

Using *n*! = Γ(*n* + 1), the probability

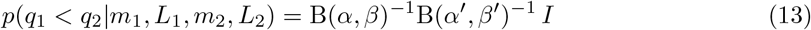

can then be written as

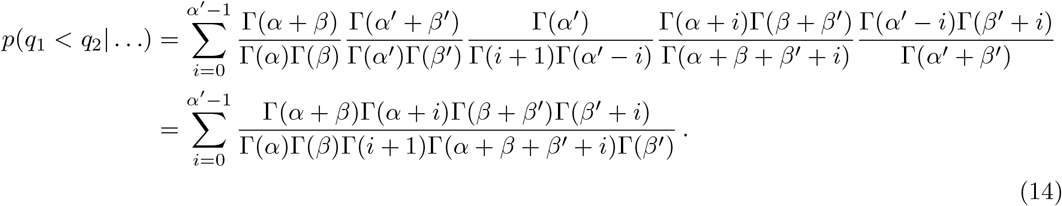

With a constant *C* and a term *π*_*i*_ depending on *i*,

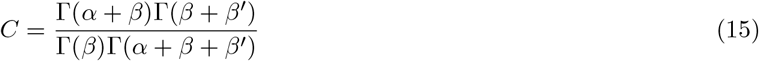

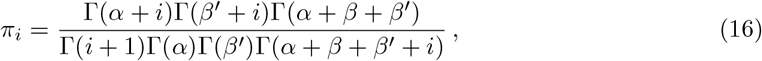

the probability 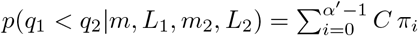 can then be computed iteratively by

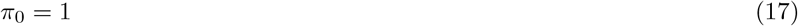

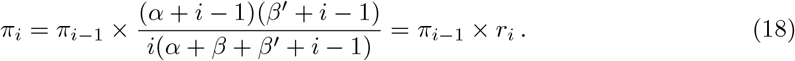

Based on the resulting probability value, the following decisions are made:

If *p*(*q*_1_ < *q*_2_|*m*_1_, *L*_1_, *m*_2_, *L*_2_) > 0.55 the (*m*_2_, *L*_2_) alignment is preferred, if *p*(*q*_1_ < *q*_2_|*m*_1_, *L*_1_, *m*_2_, *L*_2_) < 0.45 the (*m*_1_, *L*_1_) alignment is preferred. If the probability falls into the range 0.45 to 0.55 we consider it as “inconclusive”, and prefer the extension that offers the longer extension instead.

The probability *p*(*q* < *q*′|*α, β, α*′, *β*′) can be computed using the following pseudocode:

~~~
prob(a1, b1, a2, b2)
  if a1 < a2:
     return 1.0 - prob(a2, b2, a1, b1)
  sum = pi = 1.0
  for i = 1.0 to a2-1
     pi *= (a1+i-1) * (b2+i-1) / i / (a1+b1+b2+i-1)
     sum += pi
  return computeConstant(a1, b1, a2, b2) * sum
~~~

## References

[1] Sunagawa, S. et al. Tara Oceans: towards global ocean ecosystems biology. Nature Rev Microbiol 18, 428–445 (2020).

[2] Morais, L. H., Schreiber IV, H. L. & Mazmanian, S. K. The gut microbiota–brain axis in behaviour and brain disorders. Nature Rev Microbiol 19, 241–255 (2021).

[3] Round, J. L. & Mazmanian, S. K. The gut microbiota shapes intestinal immune responses during health and disease. Nature Rev Immunol 9, 313–323 (2009).

[4] Amabebe, E., Robert, F. O., Agbalalah, T. & Orubu, E. S. Microbial dysbiosis-induced obesity: role of gut microbiota in homoeostasis of energy metabolism. British J Nutrition 123, 1127–1137 (2020).

[5] Roux, S., Matthijnssens, J. & Dutilh, B. E. Metagenomics in virology. Encyc Virol 133 (2021).

[6] Adriaenssens, E. M. et al. Metagenomic analysis of the viral community in Namib Desert hypoliths. Env Microbiol 17, 480–495 (2015).

[7] Schulz, F. et al. Hidden diversity of soil giant viruses. Nature Communications 9, 1–9 (2018).

[8] Santos-Medellin, C. et al. Viromes outperform total metagenomes in revealing the spatiotemporal patterns of agricultural soil viral communities. ISME J 15, 1956–1970 (2021).

[9] Roux, S. et al. Ecogenomics and potential biogeochemical impacts of globally abundant ocean viruses. Nature 537, 689–693 (2016).

[10] Coutinho, F. H. et al. Marine viruses discovered via metagenomics shed light on viral strategies throughout the oceans. Nature Communications 8, 1–12 (2017).

[11] Hwang, J., Park, S. Y., Park, M., Lee, S. & Lee, T.-K. Seasonal dynamics and metagenomic characterization of marine viruses in Goseong Bay, Korea. PloS one 12, e0169841 (2017).

[12] Gregory, A. C. et al. Marine DNA viral macro-and microdiversity from pole to pole. Cell 177, 1109–1123 (2019).

[13] Wolf, Y. I. et al. Doubling of the known set of RNA viruses by metagenomic analysis of an aquatic virome. Nature Microbiol 5, 1262–1270 (2020).

[14] Zayed, A. A. et al. Cryptic and abundant marine viruses at the evolutionary origins of earth’s rna virome. Science 376, 156–162 (2022).

[15] Shkoporov, A. N. et al. The human gut virome is highly diverse, stable, and individual specific. Cell host & microbe 26, 527–541 (2019).

[16] Gregory, A. C. et al. The gut virome database reveals age-dependent patterns of virome diversity in the human gut. Cell host & microbe 28, 724–740 (2020).

[17] Gulyaeva, A. et al. Discovery, diversity, and functional associations of crAss-like phages in human gut metagenomes from four Dutch cohorts. Cell reports 38, 110204 (2022).

[18] Li, R., Wang, Y., Hu, H., Tan, Y. & Ma, Y. Metagenomic analysis reveals unexplored diversity of archaeal virome in the human gut. Nature Communications 13, 7978 (2022).

[19] Sutton, T. D. & Hill, C. Gut bacteriophage: current understanding and challenges. Front Endocrinol 10, 490764 (2019).

[20] Breitbart, M., Bonnain, C., Malki, K. & Sawaya, N. A. Phage puppet masters of the marine microbial realm. Nature microbiology 3, 754–766 (2018).

[21] Koonin, E. V., Krupovic, M. & Dolja, V. V. The global virome: How much diversity and how many independent origins? (2023).

[22] Nayfach, S. et al. Metagenomic compendium of 189,680 DNA viruses from the human gut microbiome. Nature Microbiology 6, 960–970 (2021).

[23] Mirzaei, M. K. et al. Challenges of studying the human virome–relevant emerging technologies. Trends in Microbiology 29, 171–181 (2021).

[24] Kleiner, M., Hooper, L. V. & Duerkop, B. A. Evaluation of methods to purify virus-like particles for metagenomic sequencing of intestinal viromes. BMC Genomics 16, 1–15 (2015).

[25] Sanjuan, R. & Domingo-Calap, P. Mechanisms of viral mutation. Cellular and molecular life sciences 73, 4433–4448 (2016).

[26] Kupczok, A., Bailey, Z. M., Refardt, D. & Wendling, C. C. Co-transfer of functionally interdependent genes contributes to genome mosaicism in lambdoid phages. Microbial Genomics 8, 000915 (2022).

[27] Schatz, M. C., Delcher, A. L. & Salzberg, S. L. Assembly of large genomes using second-generation sequencing. Genome Res 20, 1165–1173 (2010).

[28] Steinegger, M., Mirdita, M. & Soding, J. Protein-level assembly increases protein sequence recovery from metagenomic samples manyfold. Nature Methods 16, 603–606 (2019).

[29] Li, D., Liu, C.-M., Luo, R., Sadakane, K. & Lam, T.-W. MEGAHIT: an ultra-fast single-node solution for large and complex metagenomics assembly via succinct de Bruijn graph. Bioinformatics 31, 1674–1676 (2015).

[30] Nurk, S., Meleshko, D., Korobeynikov, A. & Pevzner, P. A. metaSPAdes: a new versatile metagenomic assembler. Genome Res 27, 824–834 (2017).

[31] Bushmanova, E., Antipov, D., Lapidus, A. & Prjibelski, A. D. rnaSPAdes: a de novo transcriptome assembler and its application to RNA-Seq data. GigaScience 8, giz100 (2019).

[32] Sutton, T. D., Clooney, A. G., Ryan, F. J., Ross, R. P. & Hill, C. Choice of assembly software has a critical impact on virome characterisation. Microbiome 7, 1–15 (2019).

[33] Antipov, D., Raiko, M., Lapidus, A. & Pevzner, P. A. Metaviral SPAdes: assembly of viruses from metagenomic data. Bioinformatics 36, 4126–4129 (2020).

[34] Meleshko, D., Hajirasouliha, I. & Korobeynikov, A. coronaSPAdes: from biosynthetic gene clusters to RNA viral assemblies. Bioinformatics 38, 1–8 (2022).

[35] Mallawaarachchi, V. et al. Phables: from fragmented assemblies to high-quality bacteriophage genomes. Bioinformatics 39, btad586 (2023).

[36] Fritz, A. et al. Haploflow: Strain-resolved de novo assembly of viral genomes. Genome Biology 22, 1–19 (2021).

[37] Fitzpatrick, A. H., Rupnik, A., O’Shea, H. & Cotter, P. High throughput sequencing for the detection and characterization of RNA viruses. Front Microbiol 12, 621719 (2021).

[38] Baaijens, J. A., El Aabidine, A. Z., Rivals, E. & Schönhuth, A. De novo assembly of viral quasispecies using overlap graphs. Genome Res 27, 835–848 (2017).

[39] Li, W. et al. Vipra-haplo: de novo reconstruction of viral populations using paired end sequencing data. IEEE/ACM Transactions on Computational Biology and Bioinformatics (2024).

[40] Hunt, M. et al. IVA: accurate de novo assembly of RNA virus genomes. Bioinformatics 31, 2374–2376 (2015).

[41] Yang, X. et al. De novo assembly of highly diverse viral populations. BMC Genomics 13, 1–13 (2012).

[42] Yuan, C., Lei, J., Cole, J. & Sun, Y. Reconstructing 16S rRNA genes in metagenomic data. Bioinformatics 31, i35–i43 (2015).

[43] Miller, C. S., Baker, B. J., Thomas, B. C., Singer, S. W. & Banfield, J. F. EMIRGE: reconstruction of full-length ribosomal genes from microbial community short read sequencing data. Genome Biol 12, 1–14 (2011).

[44] Vollmers, J., Wiegand, S. & Kaster, A.-K. Comparing and Evaluating Metagenome Assembly Tools from a Microbiologist’s Perspective - Not Only Size Matters! PLoS ONE 12 (2017).

[45] Steinegger, M. & Soding, J. Clustering huge protein sequence sets in linear time. Nature Communications 9, 1–8 (2018).

[46] Mikheenko, A., Saveliev, V. & Gurevich, A. MetaQUAST: evaluation of metagenome assemblies. Bioinformatics 32, 1088–1090 (2016).

[47] Craigie, R. & Bushman, F. D. HIV DNA integration. CSH Perspective Med 2, a006890 (2012).

[48] Maldarelli, F. et al. The role of HIV integration in viral persistence: no more whistling past the proviral graveyard. J Clin Invest 126, 438–447 (2016).

[49] Steinegger, M. & Soding, J. MMseqs2 enables sensitive protein sequence searching for the analysis of massive data sets. Nature Biotechnol 35, 1026–1028 (2017).

[50] Callanan, J. et al. Expansion of known ssRNA phage genomes: from tens to over a thousand. Science Adv 6, eaay5981 (2020).

[51] Wolf, Y. I. et al. Origins and evolution of the global RNA virome. MBio 9, e02329–18 (2018).

[52] Tars, K. ssRNA Phages: Life Cycle, Structure and Applications. In Biocommunication of Phages, 261–292 (Springer, 2020).

[53] Chamakura, K. R. et al. Rapid de novo evolution of lysis genes in single-stranded RNA phages. Nature Communications 11, 1–11 (2020).

[54] Quast, C. et al. The SILVA ribosomal RNA gene database project: improved data processing and web-based tools. Nucleic Acids Res 41, D590–D596 (2013).

[55] Edgar, R. C. Updating the 97% identity threshold for 16S ribosomal RNA OTUs. Bioinformatics 34, 2371–2375 (2018).

[56] Deng, Z.-L. et al. Evaluating assembly and variant calling software for strain-resolved analysis of large DNA viruses. Brief Bioinformatics 22, bbaa123 (2021).

[57] Bouras, G. et al. How low can you go? Short-read polishing of Oxford Nanopore bacterial genome assemblies. Microbial Genomics 10, 001254 (2024).

[58] Hall, M. B. et al. Benchmarking reveals superiority of deep learning variant callers on bacterial nanopore sequence data. bioRxiv 2024–03 (2024).

[59] Lauber, C. & Seitz, S. Opportunities and challenges of data-driven virus discovery. Biomolecules 12, 1073 (2022).

[60] Koonin, E. V. & Yutin, N. The crass-like phage group: how metagenomics reshaped the human virome. Trends in Microbiology 28, 349–359 (2020).

[61] Benler, S. et al. Thousands of previously unknown phages discovered in whole-community human gut metagenomes. Microbiome 9, 1–17 (2021).

[62] O’Leary, N. A. et al. Reference sequence (RefSeq) database at NCBI: current status, taxonomic expansion, and functional annotation. Nucleic Acids Res 44, D733–D745 (2016).

[63] Sayers, E. W. et al. GenBank. Nucleic Acids Res 47, D94–D99 (2019).

[64] Bushnell, B. BBMap: a fast, accurate, splice-aware aligner. Tech. Rep., Lawrence Berkeley National Lab.(LBNL), Berkeley, CA (United States) (2014).

[65] Sczyrba, A. et al. Critical assessment of metagenome interpretation—a benchmark of metagenomics software. Nature Methods 14, 1063–1071 (2017).

[66] Meyer, F. et al. Critical assessment of metagenome interpretation: the second round of challenges. Nature Methods 19, 429–440 (2022).

[67] Martin, M. Cutadapt removes adapter sequences from high-throughput sequencing reads. EMBnet. journal 17, 10–12 (2011).

[68] Bolger, A. M., Lohse, M. & Usadel, B. Trimmomatic: a flexible trimmer for Illumina sequence data. Bioinformatics 30, 2114–2120 (2014).

[69] Hyatt, D. et al. Prodigal: prokaryotic gene recognition and translation initiation site identification. BMC Bioinfo 11, 1–11 (2010).

[70] Burge, S. W. et al. Rfam 11.0: 10 years of RNA families. Nucleic Acids Res 41, D226–D232 (2013).

## References

[1] J. Callanan, S. R. Stockdale, A. Shkoporov, L. A. Draper, R. P. Ross, and C. Hill, “Expansion of known ssRNA phage genomes: from tens to over a thousand,” Science Adv, vol. 6, no. 6, p. eaay5981, 2020.

[2] M. Steinegger, M. Mirdita, and J. Söding, “Protein-level assembly increases protein sequence recovery from metagenomic samples manyfold,” Nature Methods, vol. 16, no. 7, pp. 603–606, 2019.

